# The nanoscale anatomy of exocytic dense-core vesicles in neuroendocrine cells

**DOI:** 10.1101/2020.08.19.257733

**Authors:** Bijeta Prasai, Gideon J. Haber, Marie-Paule Strub, John A. Ciemniecki, Kem A. Sochacki, Justin W. Taraska

## Abstract

Rab-GTPases and their interacting partners are key regulators of secretory vesicle trafficking, docking, and fusion to the plasma membrane in neurons and neuroendocrine cells. Where and how these proteins are positioned and organized with respect to the vesicle and plasma membrane are unknown. Here, we use correlative super-resolution light and platinum replica electron microscopy to map Rab-GTPases (Rab27a and Rab3a) and their effectors (Granuphilin-a, Rabphilin3a, and Rim2) at the nanoscale in 2D. Next, we develop a targetable genetically-encoded electron microscopy labeling method that uses histidine based affinity-tags and metal-binding gold-nanoparticles to determine the axial location of exocytic proteins using electron tomography. Our data show that Rab-GTPases and their effectors are distributed across the entire surface of individual docked vesicles. This circumferential distribution likely aids in the efficient transport, capture, docking, and rapid fusion of vesicles in excitable cells. The nanoscale molecular model of dense core vesicles generated from our methods reveals how key proteins assemble at the plasma membrane to regulate membrane trafficking and exocytosis.

## Introduction

In the cell’s cytoplasm, activated Rab proteins bind to intracellular membranous organelles and recruit effector and adaptor molecules. These modular multi-protein complexes dynamically associate with motors and t-SNAREs to coordinate vesicle movement, tethering, docking, and fusion.^1–4^ These protein assemblies are key for maintaining the directed flow of materials through the cell’s membrane trafficking system.^5–8^ Rab-GTPases and their binding partners are found in all eukaryotes. However, in neurons, endocrine, and exocrine cells, Rabs coordinate vesicle transport and the release of neurotransmitter, neuropeptide, and hormones by calcium-triggered exocytosis.^2, 5, 9^ This process is tightly regulated to ensure that exocytosis occurs with extreme spatial and temporal control.^10^

Work from the past several decades has identified the Rab-GTPases Rab27a and Rab3a and their effectors Granuphilin-a, Rabphilin3a, and Rim2 as drivers of dense core and microvesicle tethering and docking in Chromaffin, Ins-1, and PC12 cells.^11–14^ These studies were mainly focused on the physiological role of Rabs in the distinct stages of exocytosis. How these proteins assemble on individual vesicles, however, is unknown and remains one of the major gaps in the understanding of exocytosis.

In an attempt to fill this gap, two previous cornerstone studies used methods including mass spectrometry, quantitative immunoblotting, or imaging along with spatial modeling to develop a cartoon model of an exocytic vesicle and an entire synapse.^15, 16^ These models have been instrumental in guiding hypotheses for how vesicle proteins drive fusion. Yet, a direct physical picture of a single secretory vesicle generated from nanoscale imaging has been missing. Understanding this structure is key to understanding the regulation of exocytosis.

Here, we use correlative super-resolution fluorescence (dSTORM) and platinum replica electron microscopy (PREM) to directly visualize proteins associated with single secretory dense core vesicles (DCVs) in cultured neuroendocrine cells.^17, 18^ We localize the key proteins Rab27a, Rab3a, Granuphilin-a, Rabphilin3a, and Rim2 on identified DCVs, and quantify their nanometer scale positions across entire vesicle populations docked to the plasma membranes. Next, we develop, test, and use a new histidine-based genetically-encoded electron microscopy platinum replica labeling method to obtain a three dimensional (3D) view of proteins on single DCVs at 2-3 nanometer resolution.^18–20^ This method employs genetically encoded histidine tags to attach nickel-NTA-labeled gold nanoparticles (10 nm) on proteins.^21–23^ These gold labeled proteins were imaged with platinum replica EM tomography to pinpoint their 3D location with nanoscale precision. These data provide a new comprehensive view of proteins assembled in and around exocytic vesicles at the membrane of an excitable cell.

We conclude that Rabs and their effectors are distributed globally around docked vesicle membranes. This distribution could support the efficient capture of spherical vesicles moving through the cytosol. Because vesicles can dock and fuse within milliseconds,^24, 25^ this physical orientation and lack of clustering, layering, or reorganization during docking would allow for the extremely rapid attachment and near-instantaneous fusion at the plasma membrane in excitable cells.^26^

## Results

To image exocytic dense core vesicles (DCVs) we unroofed PC12 cells with a gentle sheering force. This treatment exposes the surface of a cell’s interior plasma membrane attached to the coverslip.^27^ The cytosol and untethered organelles are washed away.^28^ The plasma membrane that remains contains bound organelles including cytoskeletal filaments, membrane proteins, endocytic, exocytic vesicles, and unknown or unidentified objects.^27^ When these living membranes are rapidly fixed, stabilized, dried, and coated with a thin layer of platinum and carbon, a high-contrast image of the membrane replica can be acquired with transmission electron microscopy—a method commonly called platinum replica electron microscopy (PREM).^29–31^

Previously,^32, 33^ we used light microscopy to map dozens of DCV-related proteins^10, 12, 13, 34–36^ in living pancreatic beta INS-1 cells and neuroendocrine PC12 cells. We found that Rab proteins and their effectors have some of the highest correlation values with dense core vesicles.^32, 33^ These images, however, were obtained with diffraction limited total-internal reflection fluorescence (TIRF) microscopy and could not determine the protein’s sub-organellar positions. Thus, to map these proteins at the nanoscale we first confirmed the associations between Rabs and DCVs in unroofed PC12 cells using TIRF microscopy. We imaged two Rab-GTPases (Rab3a and Rab27a) and three effectors (Rabphilin3a, Rim2, and Granuphilin-a). We expressed mCherry or mRFP labeled Rab or Rab-effector proteins along with a specific marker for DCVs—mNeonGreen labeled Neuropeptide-Y (NPY-mNG), unroofed, fixed and imaged these membranes with TIRF microscopy. Figure 1a and b show that these proteins are highly correlated with labeled DCVs in unroofed cells. These data match measurements from both live and intact cells (Supplementary Fig. 1b). Thus, unroofing does not substantially alter the fluorescent correlation values between these proteins and vesicles.

**Figure 1.**
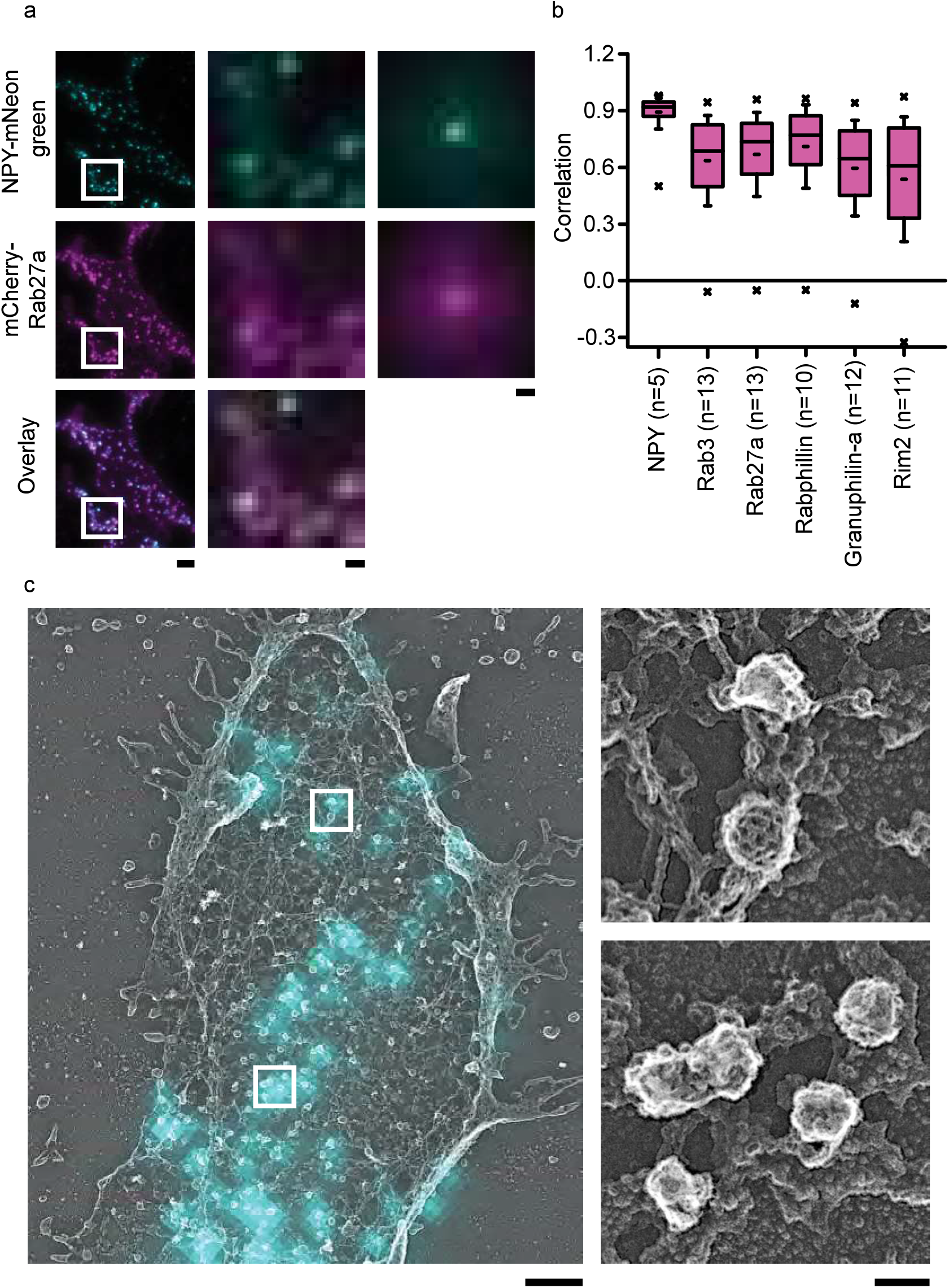
Colocalization study with TIRF and DCV visualization with PREM. (a) TIRF microscopy images of PC12 cells co-transfected with NPY-mNG (first row), Rab27a-mCherry (second row), and their overlay (third row). Left column shows representative cell image with scale bar, 3 μm. Middle column shows the enlarged images from white boxed regions. Scale bar is 0.5 μm. Small regions from 5 cells normalized to the brightest pixel and averaged together (Right column, Scale bar is 0.5 μm). (b) Correlation analysis of 5 proteins with NPY-GFP-labeled DCVs in PC12 cells. Cells are sorted based on their mean correlation values. Pink boxes are the 25th-75th percentile range of data, and the whiskers are the SD. The solid bar is the median, and the small dash is the mean. The × marks above and below each data set are the 1st and 99th percentiles, (c) TIRF NPY-mNG image (cyan) overlaid with PREM image (grey) of the unroofed PC12 cell membrane. Single DCV and DCV cluster (right panel) from white boxes in the left panel. Scale bars are 1 μm and 100 nm for left and right panels, respectively.

Correlative super-resolution light and electron microscopy can determine the position of proteins in the dense structural environment of the cell at the nanoscale.^18^ Yet, to correlate fluorescence images with EM structures of known identity, objects visible in EM must be recognizable by shape, texture, or position. Dense core vesicles in platinum replica images appear as smooth round objects and are difficult to unambiguously identify solely from PREM images. Specifically, in unroofed PC12 cells, DCVs appear as smooth spheres that are randomly scattered across the membrane (Supplementary Fig. 1d, highlighted blue). Here, we could mistakenly assign fluorescence to an organelle that is not a bona fide DCV. As a counterexample, clathrin coated vesicles have a unique honeycomb lattices that are easy to identify (Supplementary Fig. 1d, highlighted yellow). To overcome this challenge, we developed a new tripartite TIRF, super-resolution localization, and PREM-based correlative imaging pipeline to mark DCVs. This allowed for later super-resolution images to be assigned to validated DCV structures in platinum replica images.

To mark vesicles and identify DCVs, we used the fact that DCVs are loaded with NPY and completely release this soluble cargo when triggered to undergo exocytosis with depolarization.^37^ Thus, a smooth round vesicle seen in PREM that contains an NPY-mNG fluorescence signal in the correlative image is a DCV attached to the plasma membrane that has not yet fused. We first obtained TIRF images of unroofed PC12 cells expressing NPY-mNG. These diffraction-limited NPY-mNG fluorescence spots were then assigned as vesicles, marked, and further studied (Fig. 1c). Next, we used our previously developed correlative dSTORM and PREM CLEM method^18^ to map the location of super-resolution fluorescence signals with respect to these TIRF-validated DCVs. This allowed us to perform a precise 2D nanoscale colocalization at identified and verified DCVs. To image proteins, proteins of interest were fused with non-fluorescent dark GFP proteins, expressed, and labeled with Alexa Fluor 647 conjugated GFP nanobodies. Cells were also labeled with Phalloidin 568 to highlight the cell’s shape. Last, cells of interest were prepared for EM, replicas were made, and the images of these three modalities were digitally correlated to align the nanoscale protein localization within the cellular context on verified dense core vesicles (Supplementary Fig. 2d-f).

Figure 2a-e shows dSTORM images aligned with PREM images for proteins labeled with Alexa Fluor 647 conjugated GFP nanotrap. CLEM images for all five proteins (Rab3a, Rab27a, Rabphilin3a, Rim2, and Granuphilin-a) show fluorescence on or close to dense core vesicles. To measure the association of these proteins with vesicles, we generated fluorescence profiles by outlining the NPY-mNG identified vesicles in EM and analyzing the average normalized fluorescence pixel values as a function of distance from either the center or edge of the vesicle. The detailed analysis workflow is shown in Supplementary Fig. 2. For radial profiles (Fig. 2h), we binned pixels in 12 nm increments from the center of a vesicle up to 18 bins (Fig. 2f-g). And, for edge profiles (Supplementary Fig. 2k), we binned as previously described for clathrin coated pits^17^ —5 bins inward and 10 bins away from the vesicle edge. Both radial and edge profiles show distinctive spatial distributions of Rab and Rab effector proteins on DCVs. The vesicle size analysis showed that the DCV radii ranged from 60.7±10.2 nm to 70±12 nm for Rabphilin-3a and Rab27a expressed cells, respectively (Supplementary Fig. 1c, Supplementary Table 1). The steep fluorescence observed within this range demonstrates that these exocytic proteins are predominantly located directly on DCVs. The fluorescence profiles generated for immunolabeled endogenous proteins (Rab3a, Granuphilin-a, and Rim2) exhibited a similar fluorescence distribution as the transiently expressed fusion proteins. Specifically, both radial and edge profiles show that the endogenous Rab3a, Granuphilin-a and Rim2 are located on DCV structures (Supplementary Fig. 2l, m). Interestingly, endogenous Rim2 showed a slightly different profile compared to the former two proteins and seemed to be located closer to the edge rather than the center of the vesicle. Vesicle sizes were similar among immunolabeled PC12 cells (59.5±10.9 nm for Anti-Rab3a to 69.7±11.7 nm for Anti-Granuphilin-a) compared with fusion proteins expressed cells (60.9±10.0 nm for Granuphilin-a to 64.8±10.2 nm for Rab3a) (Supplementary Fig. 1c). Expression of GFP-tagged proteins did not substantially alter the distribution of these proteins across the vesicle population. Taken together, our results determine the distribution of key exocytic proteins on DCVs at the nanometer scale. All proteins appeared to have similar global distributions without obvious structural heterogeneity at the nanoscale. Specifically, we could detect no ring-like, or biased organization of exocytic proteins similar to that seen for components of clathrin coated pits.^17^ From these data, however, it was still unclear if exocytic proteins have layered positions relative to the plasma membrane in the vertical dimension. Thus, we developed a new method to obtain a detailed three-dimensional picture of a single DCV.

**Figure 2.**
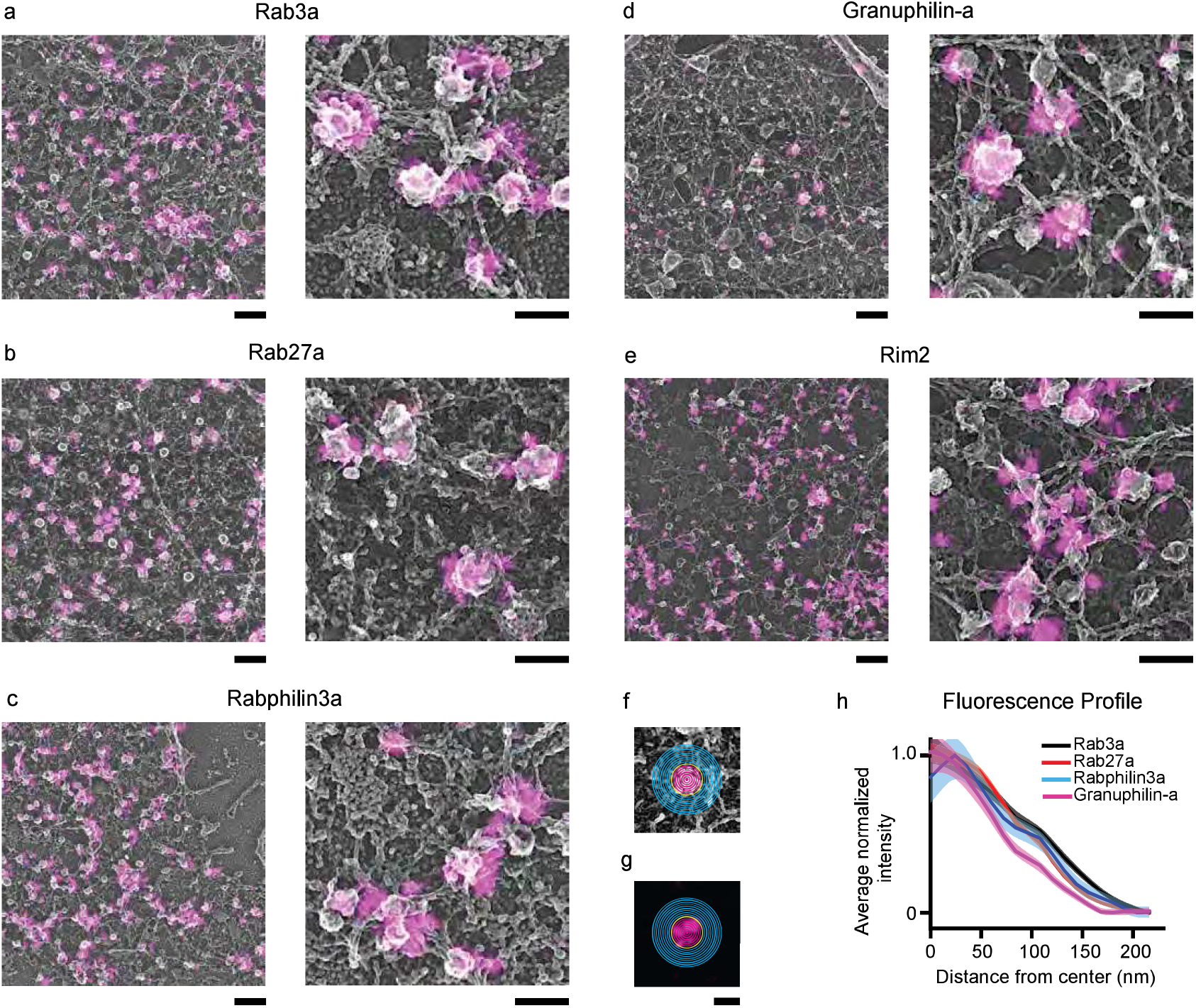
Correlative study of DCV associated exocytic proteins using dSTORM and platinum replica EM. Correlative images for PC12 cells co-transfected with NPY-mNG and AlexaFluor-647-GFP-nanobody labeled dark GFP fused proteins (a) Rab3a, (b) Rab27a, (c) Rabphilin3a, and (d) Granuphilin-a. (e) Correlative image for endogenous Rim2 protein labeled with anti-Rim2 primary antibody. Scale bars are 500 nm and 200 nm for left and right panels, respectively. These images were cropped from large images shown in Supplementary Fig. 6. The ten-pixel sized bins created from the center of the (f) EM image and applied to the respective (g) super-resolved fluorescent image to analyze fluorescence intensity, (h) Averaged and normalized fluorescence intensity profiles for Rab3a (n = 416), Rab27a (n = 523), Rabphilin3a (n = 635), Granuphilin-a (n = 251), and anti-Rim2 (n = 126). The n values are the total number of dense core vesicles analyzed (Supplementary Table 1). The mean fluorescence intensity is shown as a dark line, and the standard error of the data is shown in transparency. The fluorescence intensity profiles for endogenous proteins including Rab3a, Granuphilin-a, and Rim2 are presented in Supplementary Figs. 2l and m.

For direct protein detection with EM images, we transfected cells with proteins tagged with six histidine residues. These his-tagged proteins are avid targets for nickel-NTA gold (Ni-NTA-Au) (Fig. 3a and b). This labeling strategy offers several advantages: 1) a variety of proteins can be labeled with these short hexa-histidine tags enabling the study of many different proteins. This is challenging with immunolabeling due to the lack of specific and reliable antibodies. 2) Unlike primary/secondary antibodies, high affinity nickel-histidine interactions place gold nanoparticles close to the target (<5 nm) enabling localization with nanoscale precision.^21^ 3) Gold particles have high contrast to electron beams relative to the cellular background and platinum, making them an excellent EM markers.^38^

**Figure 3.**
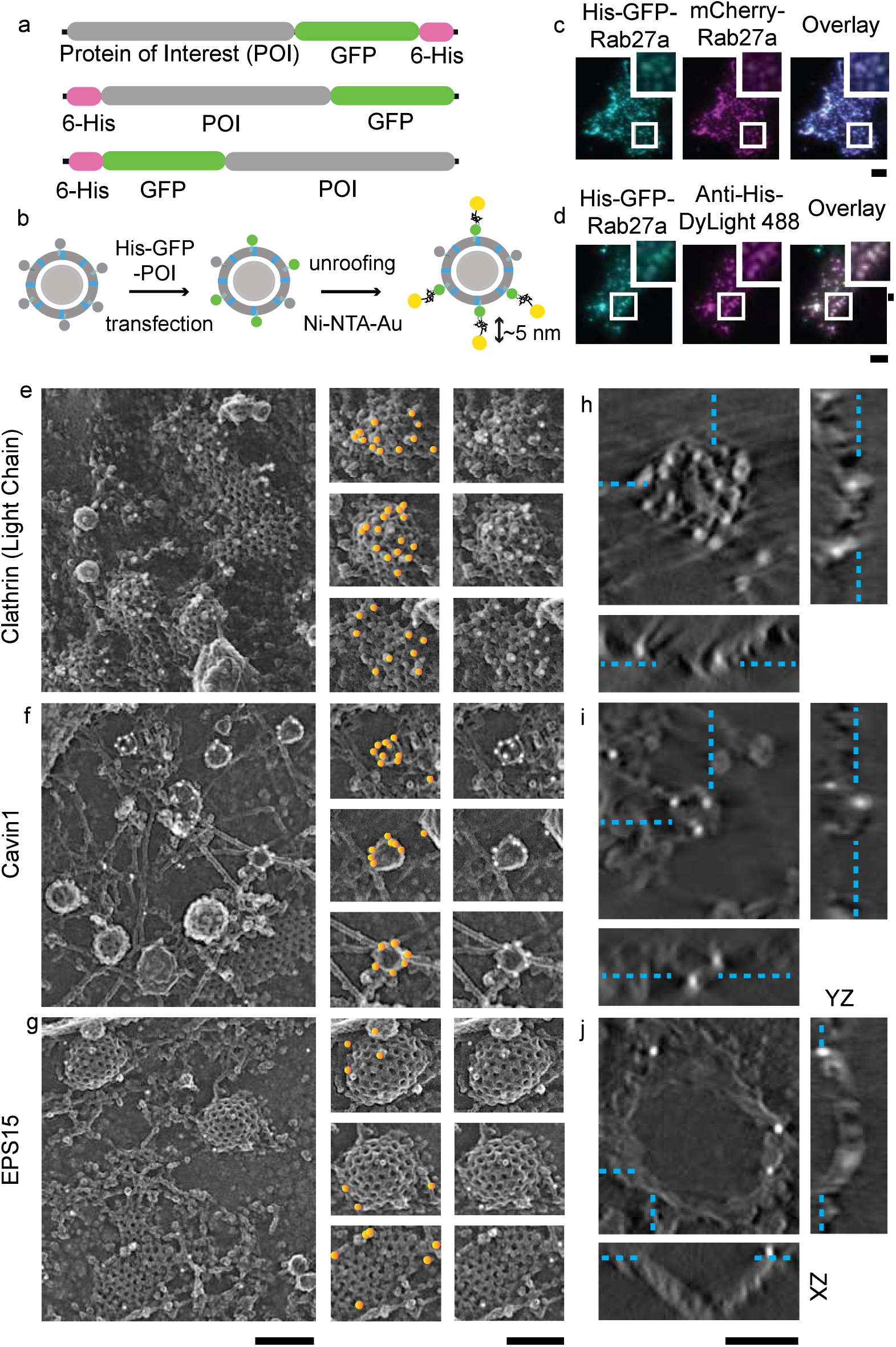
Nanogold-based protein labeling system and PREM imaging. (a) A schematic representation of plasmids used in this study with histidine and GFP fused at N or C terminal domains, (b) Scheme showing dense core vesicle associated protein labeled with Ni-NTA-Au after transfecting cells with a plasmid of interest and unroofing, (c) His-GFP-Rab27a, mCherry-Rab27a, and overlay images showing the expression of the his-tagged protein in DCVs. (d) Colocalization of His-GFP-Rab27a and Anti-Histidine-DyLight488 showing accessibility of histidine epitope. Scale bars are 3 μm. Enlarged small section (white box) for each image is at the top right comer. Scale bar is 0.5 μm. Platinum replica images of Hela cells expressed with (e) His-GFP-Clathrin Light Chain A, (f) His-Cavin-GFP, and (g) EPS15-GFP-His. Left panel scale bar is 500 nm. Enlarged images in the middle panel shows gold particles on endocytic structures marked with orange circles. The same enlarged images in the right panel shows the structures without orange markings. Scale bar is 200 nm. Tomogram section (XY view) of an individual clathrin structure labeled with Ni-NTA-Au for (h) clathrin light chain A, (i) cavin1, and (j) EPS15, and the orthogonal views in XZ and YZ dimensions. Scale bar is 200 nm. Cyan dashed lines mark the gold particles seen in XY view of a z slice and denote its location in orthogonal views.

To test this new probe system, we first evaluated the expression and localization of his-tagged proteins on DCVs as well as their suitability to labeling. Figure 3c shows the colocalization of His-tagged GFP-Rab27a with mCherry-Rab27a. Likewise, His-GFP-Rab27a, and anti-histidine DyLight 488 all strongly colocalize (Fig. 3d). This confirms that his-fusion proteins can be targeted to DCVs and are accessible to nickel-bound NTA-probes. Next, we chose two well-known endocytic proteins to evaluate the method in EM—clathrin light chain A, and cavin1. In both cases hexa-histidine tags were added to either the N or C terminal region of a GFP tagged fusion construct (Fig. 3a, scheme). Clathrin light chain A is a part of the polyhedral coat that drives clathrin mediated endocytosis.^39, 40^ Cavin1 belongs to the cavin family of proteins and is an architectural coat component of caveolae.^41^ For imaging, we selected cells with GFP fluorescence—a marker for expression of the hexa-histidine tagged proteins. We next prepared these samples for PREM and imaged them in EM. The detailed imaging pipeline is shown in Supplementary Fig. 3. The endocytic proteins used here as controls are known to envelop vesicles. Therefore, we expected vesicles to be encased in clusters of gold particles. In EM, we observed clathrin and caveolae structures clearly decorated with gold particles (Fig. 3e-f). This observation was replicated in the clathrin structures of U87-MG glioblastoma cell line (Supplementary Fig. 7). Fig. 3e-g shows gold particles highlighted with yellow spots in the central panel. Electron tomograms further confirmed that gold particles were distributed across the entire height of vesicles (Fig. 3h-i and Supplementary Videos 1-2). Of note, we observed minimal non-specific labeling in other organelles, plasma membrane regions, or cells without GFP fluorescence.

To further test the robustness of this method, we imaged EPS15, a protein known to be localized only to the rim or base of budding clathrin coated structures.^42^ In EPS15-His expressing cells labeled with Ni-NTA-gold, we found gold particles only at the periphery and bottom of clathrin (Fig. 3g, 3j, and Supplementary Video 3). As a final test, we imaged a second well-established edge protein FCH02.^17, 43^ Again, we observed a similar distribution of gold particles at the rim of clathrin coated structures (Supplementary Fig.7). The specific targeting of two established edge proteins at the rim of clathrin coated vesicles validated our method for nanoscale protein localization in 3D.

Next, we probed the unknown nanoscale 3D position of the five Rab-GTPases and their effector proteins on DCVs of PC12 cells. These proteins were tagged with six histidine residues at the N-termini, and after transfection, membrane sheets were treated with Ni-NTA-Au. Similar to the two endocytic coat proteins, we found both Rab27a, Rab3a, and their effector proteins Rabphilin3a, Granuphilin-a, and Rim2 targeted to vesicles. The gold particles were scattered across the entire surface of the vesicle (Fig. 4 a-e). In 3D, electron tomograms show that gold particles were distributed along the entire height of DCVs, similar to the positions observed for the control endocytic coat proteins (Fig. 4f-j and Supplementary Videos 4-8). For clathrin, cavin1, Rab and Rab effector proteins, we observed a range of gold particles associated with individual vesicles from as low as 2 to as many as 42 particles. For EPS15—consistent with a previously observed sparse distribution at the rim of clathrin coated vesicles and a reduced concentration in domed vesicles^17^—we found individual structures labeled with a low density of gold (1 to 7 particles).

**Figure 4.**
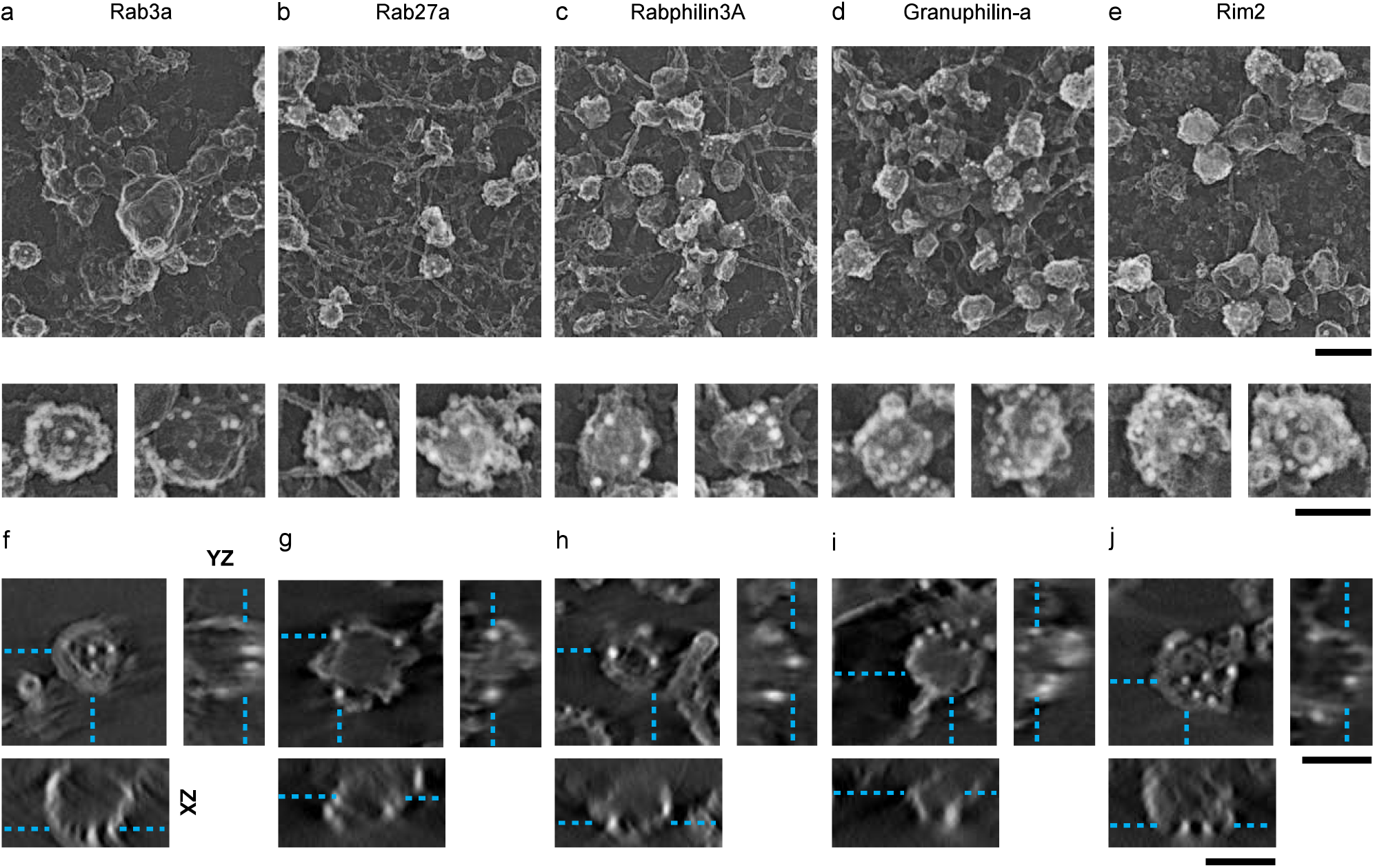
Gold nanoprobe labeled DCV proteins viewed with 2D and 3D PREM. 2D PREM images of PC12 cells transfected with His tagged (a) Rab3a, (b) Rab27a, (c) Rabphilin3a, (d) Granuphilin-a, and (e) Rim2. The upper panel shows a representative crop from larger PREM image of a cell (Supplementary Fig. 7) and lower panel shows enlarged examples of individual DCV structures labeled with Ni-NTA-Au. Scale bars are 500 nm and 200 nm, respectively. Tomogram slice (XY view) of an individual DCV structure labeled with Ni-NTA-Au for (f) Rab3a, (g) Rab27a, (h) Rabphilin3a, (i) Granuphilin-a, and (j) Rim2, and the XZ and YZ views for the z slice denoted by cyan dashed lines. Scale bar is 200 nm.

After 3D-reconstruction of electron tomograms for each protein, we analyzed the radial and axial position of gold particles with respect to the vesicle membrane. We manually outlined the sections of tomograms with a closed contour (magenta). This represents a collection of coordinate points marking the vesicle membrane. Likewise, we used points (blue) as objects to mark independent gold particles detected near these segment membranes (Fig. 5a and b). The radius and height coordinates from the membrane contours and 4001 gold points were collected from a total of 482 vesicles (Supplementary Table 2). The position of each point was identified with two values: 1) the radial distance from the centroid of the transverse cross-section, and 2) the height of the point from the plasma membrane. To analyze the distribution of proteins, we generated an average vesicle membrane profile, and assessed the radial and axial positions relative to the membranes. For every transverse cross-section in a vesicle, we first collected the radial distance from the cross-section centroid for each contour point. We normalized the radii with the point with maximum radius in each cross-section and heights with maximum height of that vesicle. Next for each particle, we normalized heights with maximum vesicle height. To normalize the particle radius in a given cross-section we drew a ray out from the centroid passing through the gold particle and reaching the contour. The contour radius at this position was used to normalize the particle’s relative position. We combined the normalized heights and radii of the contour points from a range of 54 to 65 vesicles (for clathrin light chain A and EPS15, respectively), and the associated 193 to 700 gold particles (for EPS15 and clathrin light chain A, respectively) collected from two independent experiments for each protein (Supplementary Table 2). The data were arranged in ascending order of heights. Partition averages were then obtained by dividing the heights into 10 bins. Finally, average distribution profiles were obtained by plotting normalized bin height versus normalized radial distance (Fig. 5b).

**Figure 5.**
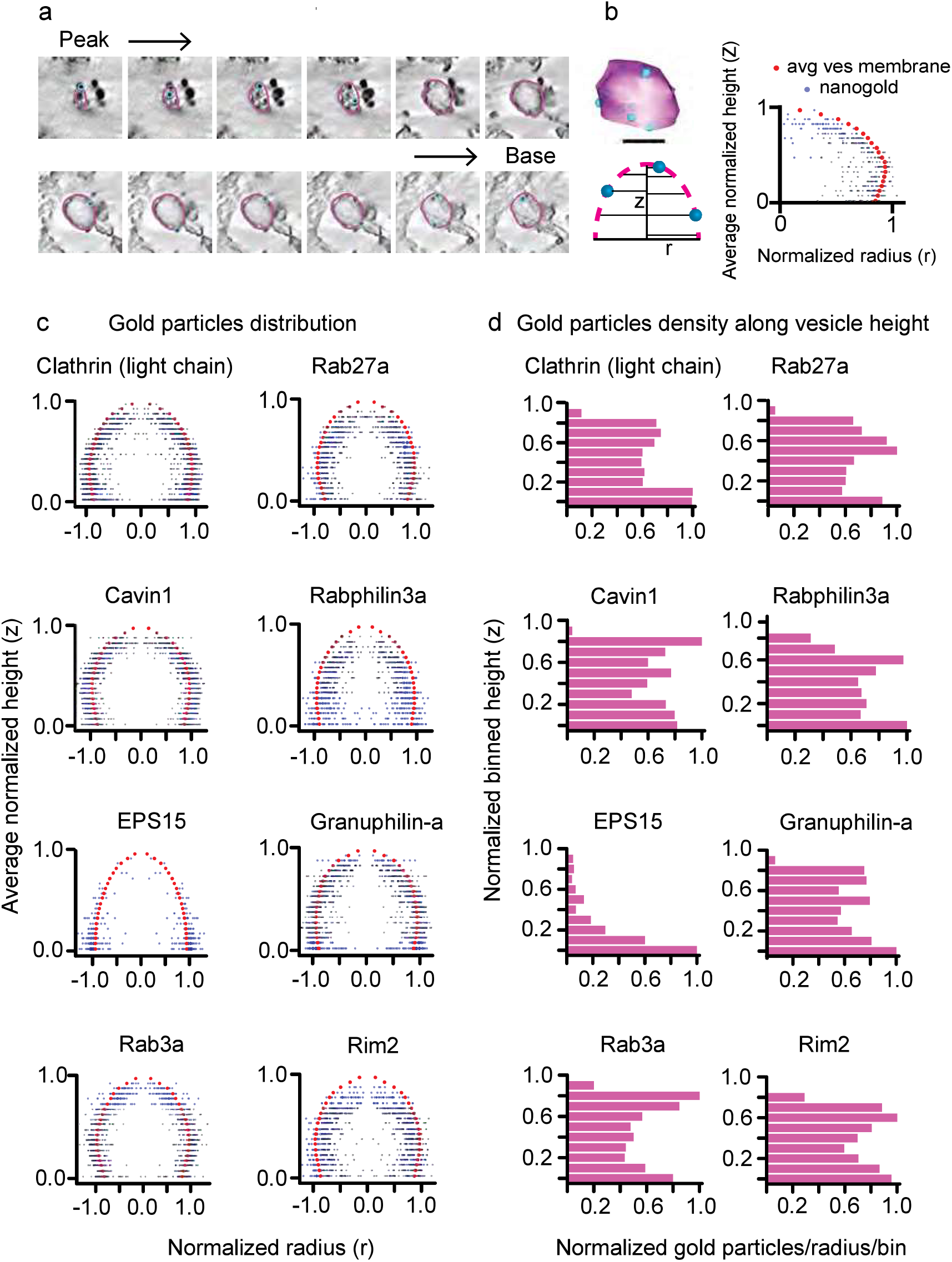
Distribution and frequency of gold particles on vesicles. (a) Tomogram z slices of a single clathrin coated vesicle from top to the base of the structure, (b) 3D model view of the vesicle from contour points for membrane vesicle and scatter points for gold particles (Upper panel, scale bar is 50 nm). Scheme (lower panel) for obtaining positions for gold particles with respect to vesicle membrane contours. Radius for each contour point (magenta, r), its height from the base of the vesicle (Z), radius for gold particle (blue), and its height. A representative profile on the right panel showing distribution of gold particles (blue) with respect to vesicle membrane (red), (c) Spatially averaged and normalized distribution profile for clathrin, cavin1, EPS15, Rab3a, Rab27a, Rabphilin3a, Granuphilin-a, and Rim2. The gold particles are plotted twice and presented symmetrically, (d) Density or frequency of gold particles representing proteins present along the vesicle height (10 bins). The histograms show the sum number of particles in bin (for all vesicles) divided by the average radius of the bin per vesicle. The number of vesicles analyzed are: clathrin light chain A, 54; Cavin1, 63; EPS15, 65; Rab3a, 59; Rab27a, 58; Rabphilin3a, 58; Granuphilin-a, 62; Rim2, 63 (Supplementary Table 2).

In Figure 5c, we show the distribution profiles of gold particles with respect to the vesicle membrane for three control endocytic and five unknown exocytic Rab proteins. Here, we plotted the same points across the ordinate to provide a visual appearance of vesicle model for visualization and interpretation. Clathrin light chain A and cavin1 plots show universal distribution of gold. This finding is in accordance with the fact that clathrin light chain A and cavin1 proteins are the framework of clathrin coated and caveolae vesicles, respectively. Further, the averaged view of gold distribution for EPS15 corroborate our EM images and proposed position of EPS15 at clathrin sites. A large number of gold particles are concentrated at the base of the model. Similar to the two endocytic coat proteins, the five DCV associated Rab proteins and their binding proteins show global distribution. Additionally, we computed the frequency of the gold particles found in the 10 separate bins to assess the uniformity of protein distribution along vesicle height. Figure 5d presents density histograms for the eight individual proteins and show normalized frequency of gold particles in partition averaged cross-section per vesicle along the vesicle height. We detected a strong difference in the distribution of the two endocytic coat proteins and the five exocytic Rab proteins and peripheral EPS15. The first seven proteins all show uniform protein density compared to EPS15. However, judging the variation in their density in a single structure is difficult. We do note that there are differences in the gold density histograms toward the top of the vesicle population. The significance of these differences cannot be substantiated because a large error in the differential cross-sectional area of the very top of the vesicles occurs in this analysis. Therefore, these slight differences are not reflected in our consensus model.

As a control we compared the results from our method with those from three-dimensional super-resolution fluorescence microscopy (3D-STORM) imaging of endogenous proteins. For this, we examined the proteins for which well-documented antibodies are commercially available including clathrin heavy chain, EPS15, Rab3a, and Granuphilin-a. We unroofed cells, immunolabeled with antibodies, and obtained 3D-STORM images according to the previously published method.^44^ Clathrin heavy chain is another component of clathrin triskelia, which cooperatively with clathrin light chain forms the polyhedral honeycomb clathrin coats.^39^ Therefore, clathrin heavy chain is expected to be localized with the same profile as clathrin light chain. 3D-STORM enables imaging with an axial resolution of 50 nm, and thus it is capable of discerning 100-200 nm sized vesicles in 3D.^44^ Images from clathrin heavy chain immunolabeled cells showed 100-200 nm domed clathrin structures (Supplementary Fig. 5). For EPS15, we did not observe domes, confirming this protein’s edge/bottom position (Supplementary Fig. 5). For Rab3a and Granuphilin-a, we saw 100-150 nm DCV foci extending from the cell membrane (Supplementary Fig. 5). With these data we conclude that our protein labeling method is effective. We further conclude that the localization of the proteins we determined was not due to overexpression. While confirmatory of our gold labeling, at this resolution, 3D-localization microscopy is insufficient to detect proteins with enough precision to formulate an accurate nanoscale structural model. Therefore, the potential of our new method to detect unexplored proteins of interests at isotropic scales approaching 1 nm in 3D is noteworthy.

## Discussion

A unified model of cell biology requires both a complete functional and structural understanding of organelles and signaling pathways.^45^ Research on the molecular identity, structure, and heterogeneity of nanoscale organelles has lagged behind due to the challenges in imaging these complex machines inside living cells and tissue. Advanced localization imaging and electron microscopy are powerful tools that enable visualization of cells at the nanoscale ^46^ Here, we have combined both to resolve the structure of docked calcium regulated dense-core secretory vesicles. Using CLEM, we analyzed the nanoscale spatial position of Rab27, Rab3a, Granuphilin-a, Rabphilin3a, and Rim2 at DCVs within the complex cellular space at the plasma membrane. Then, we developed a histidine-based genetically encoded gold probe system. This strategy enabled highly specific and robust localization of proteins associated with vesicles in a 3D ultrastructural context. We validated the method by visualizing the control coat proteins clathrin and cavin1 on clathrin coated pits and caveolae, respectively. For dense core granules, electron tomogram data and the averaged particles distribution indicated that the previously unstudied Rab proteins are located across the entire 3D surface of DCV.

Rab-GTPases are thought to impart functional identity to subcellular trafficking organelles.^47, 48^ Distinct sets of Rab-GTPases assemble and disassemble on vesicles through various steps of their life cycle from biogenesis, trafficking, docking, fusion, and endocytosis.^6^ The collective action of many accessory factors determines the precise localization and targeting of Rab-GTPases to specific organelles. These include prenylation, Rab escort protein (REFs),^49, 50^ Rab-Guanine nucleotide dissociation inhibitors (Rab-GDI),^51^ Guanine nucleotide exchange factors (GEFs),^52^ and possibly membrane receptors for Rabs.^53, 54^ These factors are likely involved in the highly specific recruitment of Rab27a and Rab3a to mature dense core vesicles. Yet, questions remain. For example, when and where do Rab-GTPases and their effectors associate with DCV membranes? How and when do Rabs bind to their effectors? How many Rab-GTPase/effectors can a single vesicle contain at any given time? Does the Rab protein organization on DCV influence the vesicle’s course of action? The answers to these questions are key to bridging the gap between the biochemistry and physiology of Rabs and their dynamic structural role in the cell.

Studies over the last few decades have led to a cartoon model of the protein/lipid complex that docks vesicles to plasma membrane.^7, 12, 55^ It commonly portrays Rab-GTPases (Rab27a or Rab3a) associated with their effector proteins (Granuphilin-a or Rabphilin3 A) exclusively at the base of a vesicle—forming a docking pedestal. Effectors are tethered to plasma membrane via SNARE proteins (Fig. 6, model I).^11, 35, 56, 57^ A concentrated docking complex on diffusing vesicles could provide mechanistic features to control exocytosis. This complex has, however, never been observed beyond the clustering of syntaxin when vesicles touch the membrane.^58–60^ A docking pedestal on the vesicle would also limit the contact area available for vesicle to dock. Our data do not support this model. In a second model, vesicles would have a more global Rab distribution when they move through the cytosol. When these vesicles dock, the proteins could then diffuse to the bottom of the vesicle (Fig. 6, model 2). In principle, globally distributed binding sites across the membrane of a moving vesicle would be favorable to efficiently capture spinning, moving vesicles at the plasma membrane. These proteins could then re-distribute to the bottom of the vesicle to concentrate docking proteins at the site of exocytosis and aid vesicle fusion. Our data again do not support this type of polarized protein assembly at the plasma membrane on vesicles. Our data support a third model of vesicle docking. Here, Rabs and their effectors are universally distributed across the entire vesicle surface rather than sequestered at a specific region—either before or after vesicle docking (Figure 6, model 3). These observations were replicated for Rab27a protein in dense core vesicles of a related insulin cell line showing its commonality to other dense core vesicle systems (Supplementary Fig.7). We did not experimentally determine when and where Rab-effector complexes form on vesicles. This could occur before or after DCVs dock. We propose that this likely occurs prior to docking as effector proteins are known to sustain Rab-GTPases in active GTP-bound form and maintain vesicle identity, interact with the cytoskeleton for regional trafficking, and bind with target membrane factors for tethering and fusion.^61, 62^ Studies have also shown that effectors have affinity for GTP-bound active Rab-GTPases.^63, 64^ This state exists only when GTPases are vesicle bound. This implies that effectors bind to Rabs that are attached to the vesicle surface. This could occur before or after the vesicle approaches the membrane. Here, in our model, we integrated our data with previous studies to suggest Rabs and recruited effectors promote DCVs to tether and dock to plasma membrane by interacting with SNARE protein complex.

**Figure 6.**
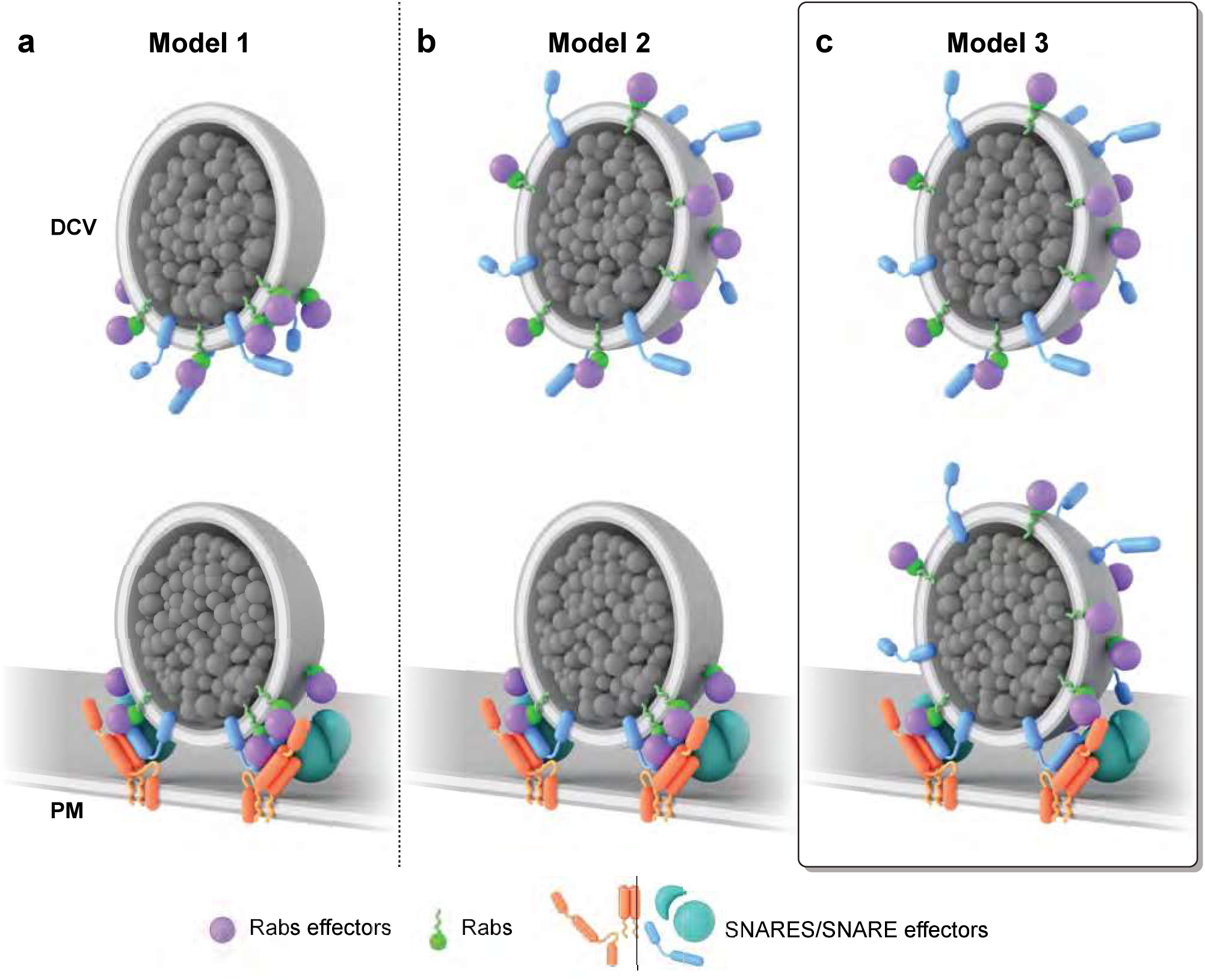
Architectural model of DCV. Scenarios for the distribution of prenylated Rab-GTPases and their respective effector proteins on a tethering and docking DCV. (a) Model 1: Illustrates the currently accepted polarized view of Rab-effector complexes on docking DCV. (b) Model 2: A hypothetical model shows the universal distribution of the protein complexes, which diffuse and polarize before tethering and docking, (c) Model 3: The model generated from our study describes the global distribution of protein complexes that enables DCVs to tether and dock at plasma membrane.

What controls vesicle docking to specific sites on the plasma membrane before fusion is still unknown. Rab-GTPases and effectors likely help tether and engage with suitable docking factors including syntaxin and MUNC18 to provide directionality for DCVs trafficking.^8, 65^ The circumferential distribution of Rabs that we observed could be a means to effectively target diffusing secretory vesicles. For example, in situations when tethered vesicles dissociate and wobble, different sets of Rab-effector complexes on the vesicle surface may aid in rapid recapturing of the organelle preventing its escape. Further research is needed to fully understand the effects the number and density of Rab-effector complexes have on tethering, docking, and fusion of DCVs. Future work investigating the mutants of the GTPases jointly with their wild-type effectors is needed.

Our methods enables accurate visualization of protein locations on single vesicles. But, at this time it lacks the ability to count the endogenous number of proteins present on a vesicle surface at a given time. Other works have quantified more than a dozen major synaptic vesicle associated proteins. One study found 10 Rab3a proteins on average associated with SVs.^15^ This was verified by another study on the composition of synaptosome.^16^ Dense core vesicles are 2-3 times larger than synaptic vesicles,^66^ but share similar docking, tethering, and fusion proteins.^67^ Immunoprecipitation assays on fractionated PC12 cells have shown that the common proteins found in synaptic-like microvesicles and DCVs differ significantly in content.^34^ Therefore, it is likely that DCVs have a higher number of Rab proteins than SVs. Our protein labeling method combined with existing biophysical, analytical, and modeling techniques will be key in creating a detailed morphological and quantitative picture of single exocytic organelles.

In conclusion, we map the nanoscale 3D location of Rab27a, Rab3a, Rabphilin3a, Granuphilin-a, and Rim2 on DCVs of PC12 cells using CLEM and a gold-based protein labeling method for EM. Our results show that plasma membrane docked exocytic vesicles contain Rab-GTPases and effectors across their entire membrane surfaces. There was no evidence for clustering or layering of proteins. These findings highlight the role global Rab-GTPases-effector complex can play on the efficient transport, anchoring, and fusion of secretory vesicles to the plasma membrane. Finally, our method offers a new means to directly detect and identify proteins associated with organelles in various stages of their life cycle. These tools reveal the structural identity of subcellular organelles and help establish a unified model of their molecular morphology—a missing link for understanding the mechanism of exocytosis.

## Methods

### Cell culture and transfection

Low-passage frozen stock of PC12-GR5 cell line—originated from Rae Nishi’s lab (OHSU)—were maintained in growth media containing DMEM (Life Technologies), 10% fetal bovine serum (Life Technologies 26140-079), and 1% vol/vol penicillin/streptomycin (Invitrogen 15070-063). HeLa cells were maintained in the phenol red-free MEM growth media supplemented with 1% vol/vol Glutamax (Life Technologies 35050-061), and 1% vol/vol sodium pyruvate (Sigma S8636-100ML) and incubated at 37 °C, with 5% C02. For CLEM, the cells were plated on gold-nanorod-embedded coverslips (hestzig.com, part no. 600-400AuF) coated with poly-L-lysine (Sigma P4832) for 20 min prior to use. Coverslips were previously cleaned in boiling RCA etch solution (5:1:1 water/30% ammonia/30% hydrogen peroxide) for 10 min, and stored in 100% ethanol. Cells were transfected with 0.7 ml of Optimem (Life Technologies 31985062), 3 μl Lipofectamine 2000 (Life Technologies 11668027), and 1 μg of plasmid to express the proteins of interest for 4 h after being introduced to the cells. Next, transfected cells were incubated in phenol-red free DMEM growth medium until ready for imaging.

### TIRF microscopy and correlation analysis

TIRF imaging and analysis were done as previously described.^33^ An IX-81 inverted fluorescence microscope with a 100x/1.45 numerical objective was used. 488 and 561 nm laser lines were used to image cells transiently transfected with DCV marker NPY-mNG and mcherry or mRFP-proteins of interests. The image was projected onto an Andor Ixon 897 EMCCD through a Dual View (Biovision Technologies) containing 525Q/50 and 605Q/55 bandpass emission filters. Yellow-green 100-nm beads were visible in both channels of the image splitter and were used to align the red and green images with a projective image transform. Images were taken at the exposure of 100 ms with 500 ms interval. Pixel size was 160 nm. All analysis was performed using custom MATLAB (MathWorks, Natick, MA) scripts and ImageJ. Correlation analysis was performed as previously described.^32^

### STORM and PREM (CLEM)

After overnight transfection, cells were rinsed in intracellular buffer (70 mM KC1, 30 mM HEPES maintained at pH 7.4 with KOH, 5 mM MgCl_2_, 3 mM EGTA), and manually unroofed with 19-gauge needle and syringe using 2% paraformaldehyde (Electron Microscopy Sciences 15710) in intracellular buffer. After unroofing, the coverslips were transferred to fresh 2% paraformaldehyde in intracellular buffer for 20 min. They were then rinsed 4× with phosphate-buffered saline (PBS) and treated with blocking buffer (3% bovine serum albumin (BSA) in PBS) for 1 h at 5 °C. The cells were then labeled with 35-45 nM Alexa Fluor 647 nanobody in 1 ml blocking buffer for 1 h at room temperature. Alexa-Fluor 647 labeled nanobody was prepared as previously described.^17^ During the last 15 min of labelling, 16.5 pmol of Alexa Fluor 568-Phalloidin (Life Technologies A12380) was added. The coverslips were then rinsed 4× with PBS, and then post-fixed with 2% paraformaldehyde in PBS for 20 min and imaged immediately or refrigerated overnight. Blinking buffer was prepared and imaging performed as previously described.^17, 68^ Localization imaging was done on a Nikon NSTORM system equipped with an Andor iXon Ultra 897 emccd. First, 15×15 (~1 mm^2^) large montage was generated for NPY-mNG (488 nm), phalloidin stained actin filaments (567 nm), and protein of interests (647 nm). And then, 30000-40000 frames were collected at 9 ms camera exposure for selected cells. Localization data were processed using Nikon Elements 4.1. The identification parameters for all PC12 correlation data were: minimum height = 100-200, maximum height = 65,535, CCD baseline = 100, minimum width = 200 nm, maximum width = 400 nm, initial fit width = 300 nm, maximum axial ratio = 1.3, maximum displacement = 1 pixel. The data were filtered assuming 0.18 photons per count with 50-300 minimum photons.

After fluorescence imaging, coverslips were stored in 2% glutaraldehyde until ready for TEM sample preparation. TEM sample preparation and imaging were performed as previously described.^17, 68^ Coverslips were moved straight from glutaraldehyde into 0.1% w/v tannic acid (freshly dissolved in water) for 20 min. They were then rinsed 4× with water and placed in 0.1% w/v uranyl acetate for 20 min. The coverslips were washed 3×, then dehydrated, critical point dried, and coated with platinum and carbon. The diamond objective marked region of interest on the coverslip was imaged with 10× phase contrast to obtain a map of the region imaged in fluorescence. The replicas were lifted and placed onto Formvar/carbon-coated 75-mesh copper TEM grids (Ted Pella 01802-F) that were freshly glow-discharged. Again, the grid was imaged with 10× phase contrast to find the same region that was originally imaged in fluorescence. Each cell of interest was located on the grid prior to EM imaging. TEM imaging was performed as previously described^18^ at 15,000× magnification (1.2 nm per pixel) using a JEOL 1400 and SerialEM freeware for montaging.^69^

### Plasmids

An existing library of GFP and His-tag fusion plasmids were sequence confirmed and identified as in Supplementary Table 3.

### Ni-NTA-gold labeling and electron tomogram

Cells were plated on poly-L-lysine coated 25-mm # 1.5 coverslips (Warner Instruments) in 6-well plate. Next, cells were transfected with His-tagged plasmids of interest for 4 h and transferred to fresh growth medium overnight. After unroofing, cells were blocked in 3% BSA/PBS solution for an hour. Sample coverslip was then transferred to a sonicated (5 min) 1:5 solution of 10 nm Ni-NTA-Nanogold (Nanoprobes 2084) in PBS and incubated for total 1 h. The plate was placed on an orbital shaker for the first 15 minutes. During the last 15 min of incubation, 16.5 pmol of Alexa Fluor 647-Phalloidin (Life Technologies A22287) was added. And, then a 15 × 15-20 × 20 large image montage covering approximately 1-1.5 mm^2^ was acquired with 488 nm and 647 nm epifluorescence excitation. A map of cells containing GFP (488 nm) and phalloidin-647 (647 nm) fluorescence was created. TEM sample preparation was done as described earlier. Phase contrast images obtained before and after lifting the platinum replica were compared to Phalloidin-based fluorescence map. From this map, GFP fluorescent cells were selected for TEM. 2D TEM was used to survey the gold tagged organelles and tomograms were collected for those cells. Single-axis tilt series (−60° to 60°, 1° increments) were collected at 8,000×. The montages were stitched together, and the tilt series were reconstructed into tomograms using IMOD software.^69, 70^

### Fluorescence profile

Previous method of vesicle binning was followed. For edge fluorescence profile, ten-pixel (12 nm) bins - 10 outside and 5 inside the edge (unless the structure was too small) of a vesicle - were created by dilating or eroding the mask of each separate structure with a ten-pixel disc. For a radial profile, 18 bins were made from the center with 12 nm increment. Only structures with fluorescence were included in the fluorescence profile analysis. The average fluorescence in each bin (sum fluorescence signal divided by pixel number) created a profile for each structure. All of the profiles from each cell and DCVs were averaged together.

### 3D models

Reconstructed tomograms were segmented using 3dmod,^70^ model editing and image display program. The 3dmod drawing tool was used to manually outline the optical sections of reconstructed vesicle tomograms with closed contours. This outline is a collection of coordinate points marking the locations of vesicle membrane in the image. Scattered points were used as separate objects to mark the independent gold particles. Coordinates from these two objects were used to obtain the normalized radius and height for each contour points from vesicle membranes and gold scattered points. The closed contours points and scattered points can be displayed in a 3d model view as shown in Fig 5b.

### Gold distribution profile

All analysis was performed using custom MATLAB (MathWorks, Natick, MA) scripts. To analyze the distribution of gold particles/proteins relative to vesicles, each point was characterized with a radial distance from the centroid of the axial cross-section and the height from the plasma membrane base. For every cross-section in a vesicle, contour points were normalized to the max radial distance and height of the vesicle. Gold points were normalized to the radial distance of the membrane contour at the given angle. For each protein, the normalized heights and radii of all vesicles, and their associated gold particles collected from two independent experiments (Supplementary Fig. 4) were combined and arranged in ascending order according to their heights. The heights were binned in 10 equal portions, such that each bin was a tenth of the normalized height of the vesicle. Vesicle membrane and gold particle distribution profiles were generated by plotting the averaged binned heights for the vesicle and gold as a function of their averaged radial distance in the corresponding bins.

### Immunolabeling for CLEM and 3D-STORM

2% PFA fixed unroofed cells were rinsed with PBS and placed into blocking buffer for 1 h. The unroofed cells were then incubated for 1 h in blocking buffer containing 2 μg/mL primary antibody. HeLa cells were labeled with mouse anti-clathrin heavy chain (Invitrogen, MAI-065) at 1:1,000 and rabbit anti-EPS15 (Cell Signaling D3K8R) at 1:200. And, PC12 cells with 1:200 rabbit anti-Rab3a (Abeam ab3335), 1:125 rabbit anti-granuphilin (Abeam ab224047), and 1:250 rabbit anti-Rim2 (Synaptic systems 140103). The samples were then labeled with the appropriate Alexa Fluor 647-labeled secondary antibodies for another 1 h. 2 μg/ml goat anti-rabbit F(ab’)2 fragment (Life Technologies A-21246) and goat anti-mouse F(ab’)2 fragment secondary antibodies were used. Finally, the samples were washed with PBS and post-fixed for 20 min before imaging.

### 3D-STORM (For supplementary figure)

Previously established method of calibration and imaging was followed for 3D-STORM experiments. 100 nm tetraspeck beads (Invitrogen T7279) in PBS were imaged first on 25-mm # 1.5 coverslips in order to adjust CFI SR HP Apochromat TIRF 100× oil objective correction collar. The purpose of this adjustment was to minimize spherical aberration. The point spread function (PSF) of the beads obtained by imaging 400 nm above and below the focal plane were qualitatively assessed and correction collar carefully adjusted until symmetrical PSF was achieved above and below the focus. Next, the ellipticity of beads were imaged by adding cylindrical lens in the imaging path before the camera. The adequately separated beads at the center of field of view were selected for running calibration. The calibration curve was generated from the ratio of PSF widths in the x and y dimension at various heights (z) from the focal plane. For 3D imaging, the same blinking buffer were used as in 2D localization microscopy. 20000-30000 frames were collected at 9 ms camera exposure using 640 nm laser excitation. For 3D-STORM, a different Nikon NSTORM system equipped with an Andor iXon Ultra 897 emccd was used. Localization data were processed using Nikon Elements 5.11. The identification parameters for data were: minimum height = 300-500, maximum height = 20,000 or 65,535, CCD baseline = 100, minimum width = 200 nm, maximum width = 700 nm, initial fit width = 300 nm, maximum axial ratio = 2.5, maximum displacement = 0-1 pixel. The data were filtered assuming 0.112 photons per count. Select small regions of the imaged cell were analyzed at a time. Raw data containing information about the molecule localization in x, y, and z was further processed in N-STORM analysis application. For final image visualization, pixel size, z slices, and gaussian rendering radius were selected as 5 nm/pixel, 20 and 20 nm, respectively.

## Code Availability

Matlab code used in this study is specific to personal lab file formatting and not created for general use. However, code is available upon request of the corresponding author.

## Data Availability

All data supporting this work are available upon reasonable request to the corresponding author.

## Acknowledgments

We thank US National Heart, Lung, and Blood Institute (NHLBI) Light microscopy core and electron microscopy core for use of instruments. We also thank Dr. Sebastian Barg (Uppsala University, Uppsala, Sweden) for donating Rim2 plasmid. We thank the Taraska lab for helpful discussions and edits. JW Taraska is supported by the Intramural Research Program of the National Heart Lung and Blood Institute, National Institutes of Health.

## Author contributions

BP performed the experiments. GJH wrote programs for CLEM fluorescence intensity profile, and gold distribution profile. KAS helped with program development and data analysis. M.-P.S. and JAC expressed and purified GFP-nanobodies and helped with other plasmid preparations. BP and JWT designed experiments. BP processed and analyzed data. BP wrote and JWT edited the manuscript.

## Materials and Correspondence

Correspondence should be addressed to Justin Taraska.

---

**Supplementary Figure 1.**
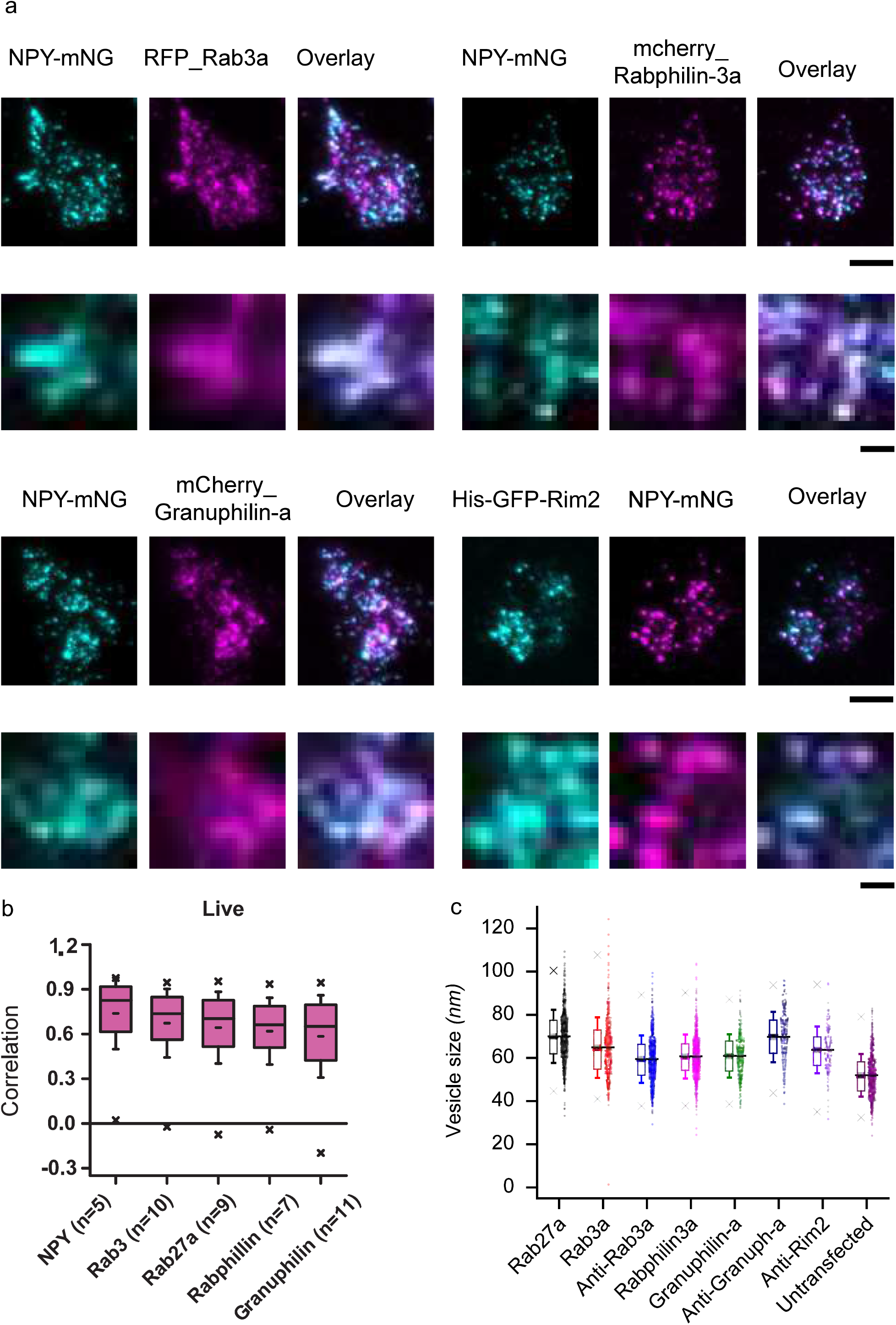
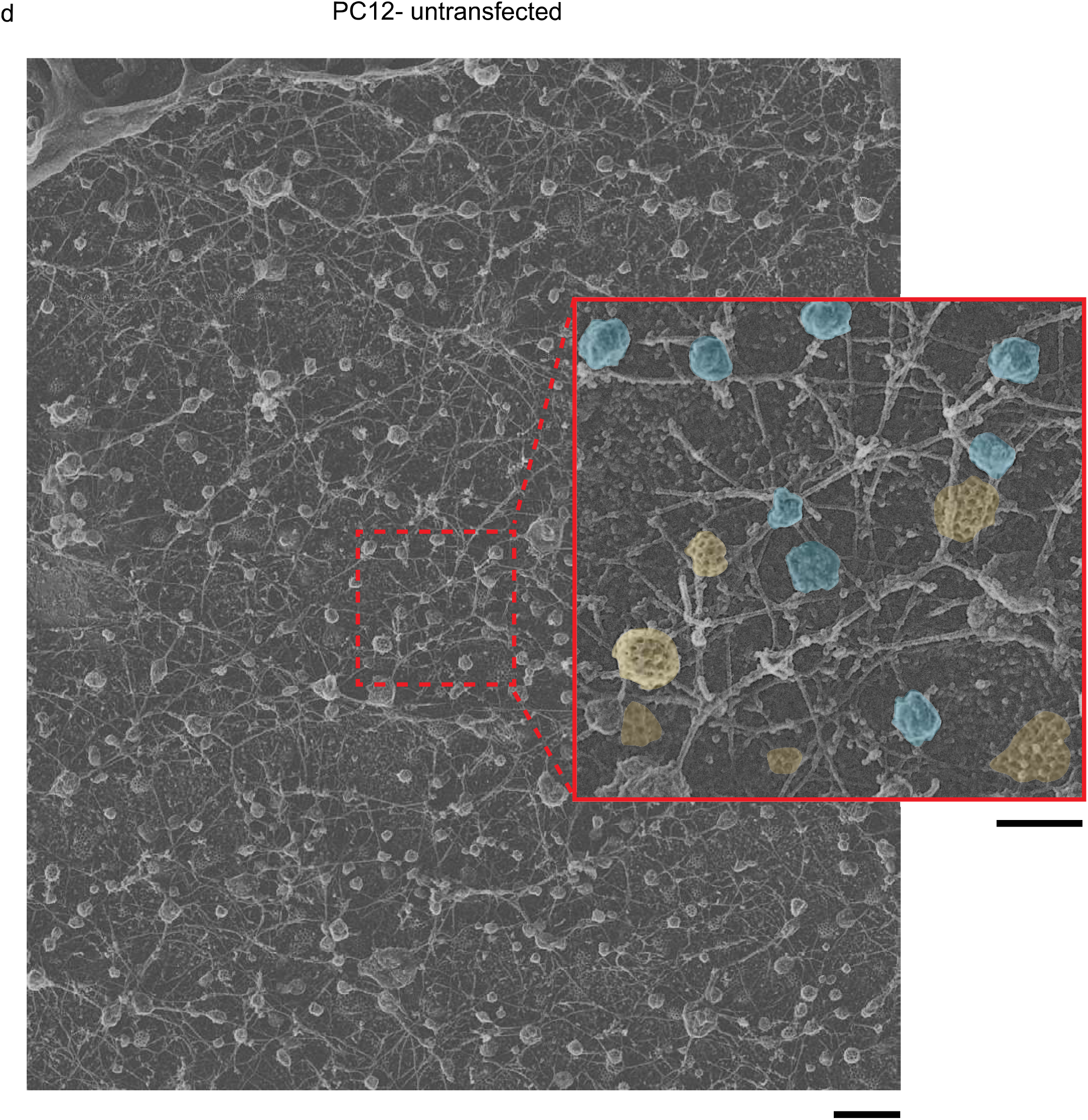
Colocalization images of unroofed cells and correlation analysis for live cells. (a) TTRF microscopy images of unroofed and fixed PC12 cells that were co-transfected with mNeongreen or mCherry labeled NPY, and mRFP or mCheriy-labeled Rab3a, Rabphilin3a, Granuphilin-a, and Rim2, and their overlay. Scale bar is 5 μm for whole cell images (upper panel) and 1 μm for zoomed in images (lower panel), (b) Correlation analysis for 5 proteins with NPY-GFP-labeled DCVs in live and intact PC12 cells. Cells are sorted based on their mean correlation values, ‘n’ denotes the number of cells used in the analysis. Boxes (magenta) are the 25th-75th percentile range of data, and the whiskers are the SD. The solid bar is the median, and the small dash is the mean. The × marks above and below each data set are the 1st and 99th percentiles, (c) Two-dimensional radius of the dense core vesicles measured on PREM images collected for unexpressed and DCV associated proteins expressed PC12 cells. Boxes are the 25th-75th percentile range of data, and the whiskers are the SD. The solid bar is the median, and the small dash is the mean. The × marks above and below each data set are the 1st and 99th percentiles. The number of vesicles used in the radius measurement: Rab3a (n = 415), Rab27a (n = 527), Rabphilin3a (n = 624), and Granuphilin-a (n = 266), and non-transfected cells (n = 315). (d) PREM image of a non-transfected PC12 cell. Scale bar is 500 nm. Enlarged image from red dashed-box shows the difference in the morphology of clathrin coated structures highlighted in yellow and DCVs highlighted in blue. Scale bar is 200 nm.

**Supplementary Figure 2.**
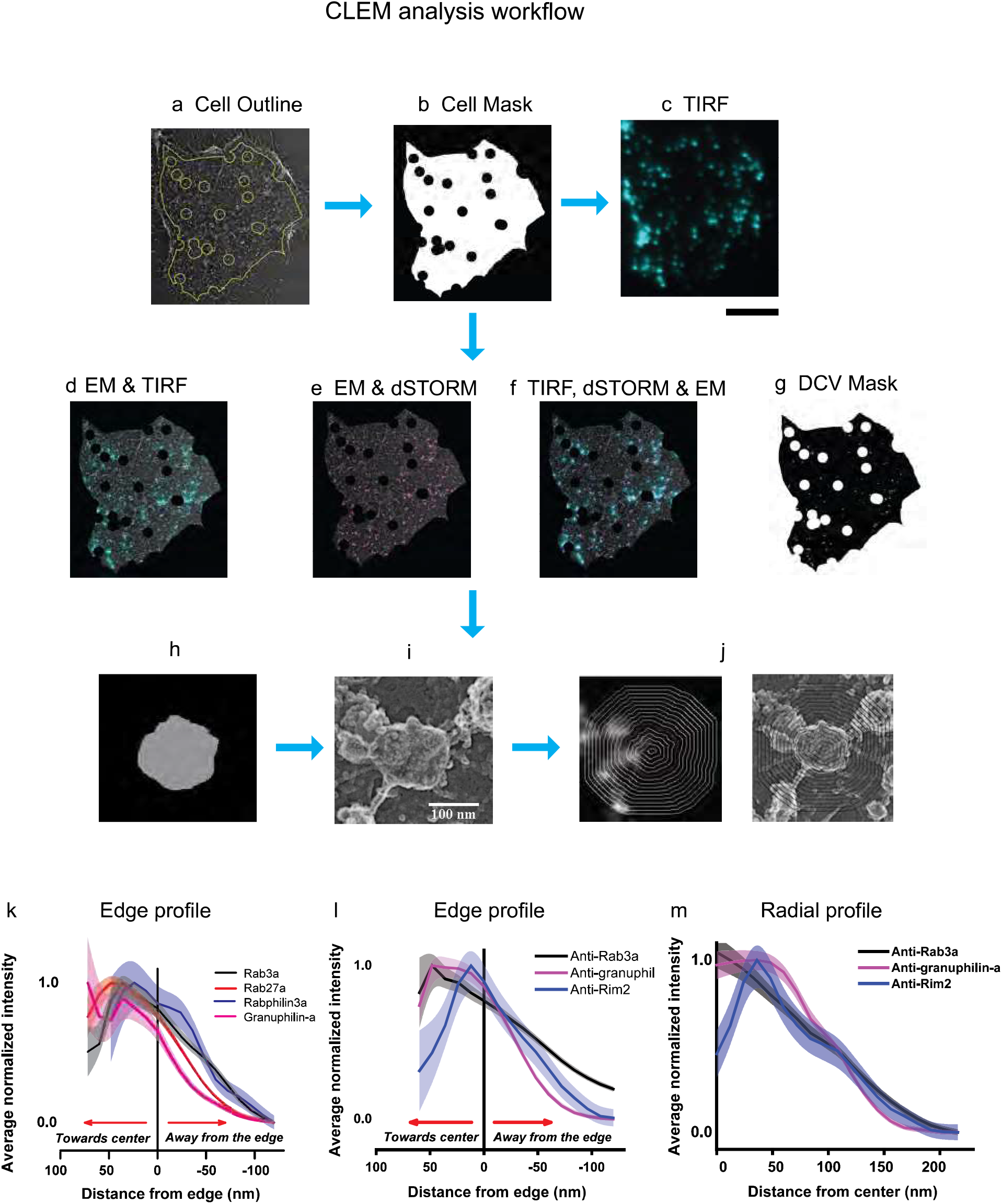
CLEM analysis workflow. (a) PREM cell membrane and gold fiducials were outlined, (b) Next, binary cell mask was created that exclude gold fiducials (~1 μm). Cell binary mask was added with (c) TIRF image to obtain (Scale bar = 5 ⊠m) (d) TIRF/EM overlay. Similarly (e) EM/dSTORM or (f) TIRF/dSTORM/EM overlay were created. Dense core vesicles were identified and outlined from either d/e/f and (g) a mask for dense core vesicles generated. Individual vesicles in (h) dSTORM and (i) EM were isolated and (j) segmented from either center or edge, (k) Edge fluorescence profiles showing normalized average fluorescence intensity distribution towards the center and away from the edge of the vesicles for cells expressed with dark GFP fused proteins. The number of vesicles used in the analysis were obtained from more than 3 cells: Rab3a (n = 415), Rab27a (n = 527), Rabphilin3a (n = 624), and Granuphilin-a (n = 277). (1) Radial fluorescence profile (m) and edge fluorescence profile for immunolabeled endogenous Rab3a, Rabphilin3a, and Rim2. The number of cells and vesicles analyzed are available in Supplementary Table 1. The structures without any fluorescence were excluded from the analysis. The standard error is shown in transparency.

**Supplementary Figure 3.**
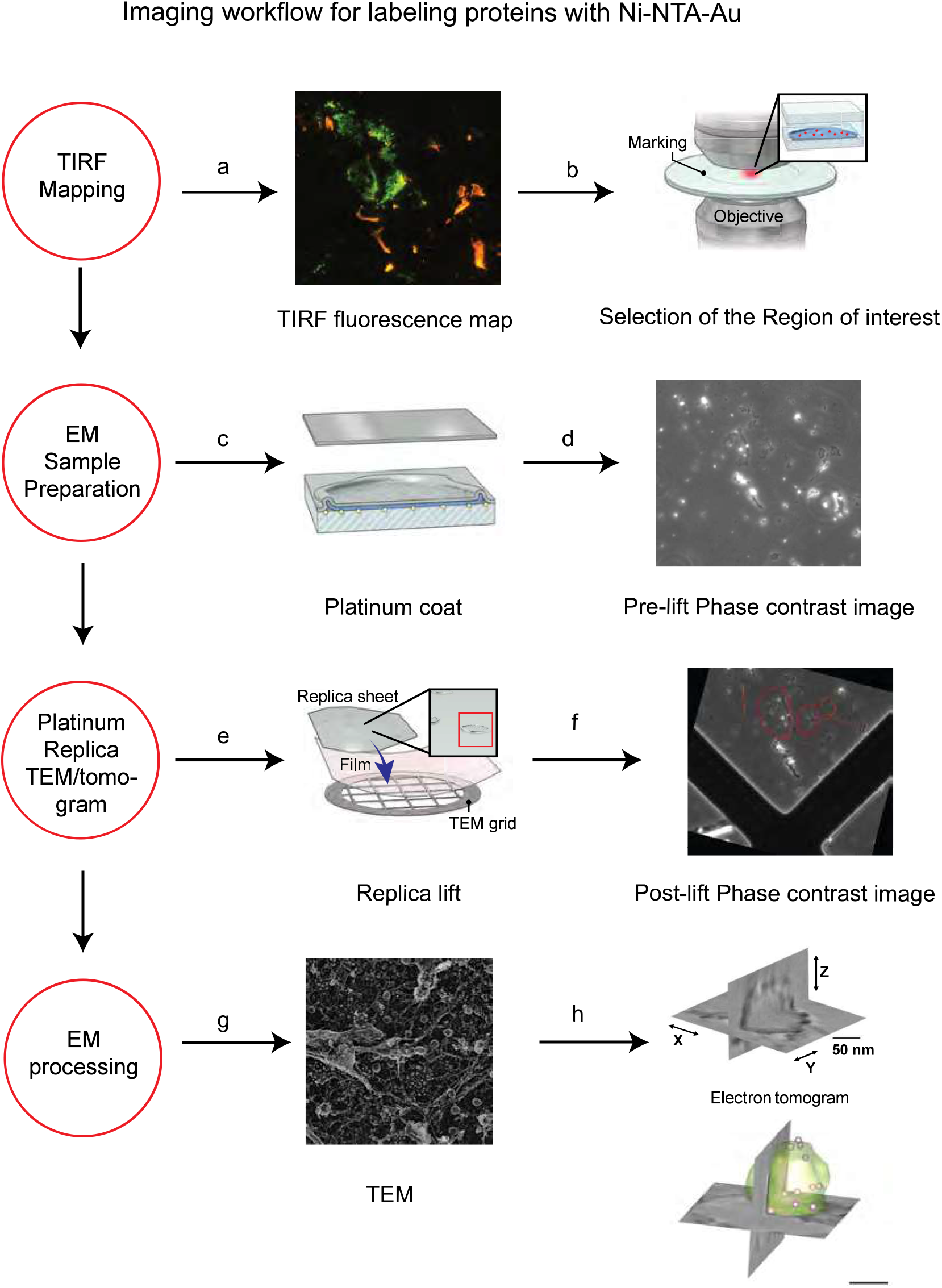
Ni-NTA-Au labeled proteins imaging and analysis pipeline. (a) Cells labeled with Ni-NTA-Au were mapped (fluorescence) to obtain a 20 × 20 large montage, (b) The mapped region was marked by etching the coverslip using a diamond objective marker, (c) The sample was then prepped for EM, critical point dried, and coated with platinum and carbon, (d) The region of interest on the coverslip was imaged with 10 × phase contrast, (e) The replica was lifted, transferred to TEM grid, and (f) imaged post-lift with phase contrast, (g) 2D TEM and (h) electron tomogram of the GFP fluorescent cells acquired with single-axis tilt series (−60° to 60°, at 1° increment) using JEOL 1400 TEM and tomograms were reconstructed and processed with IMOD software. Scale bar is 50 nm.

**Supplementary Figure 4.**
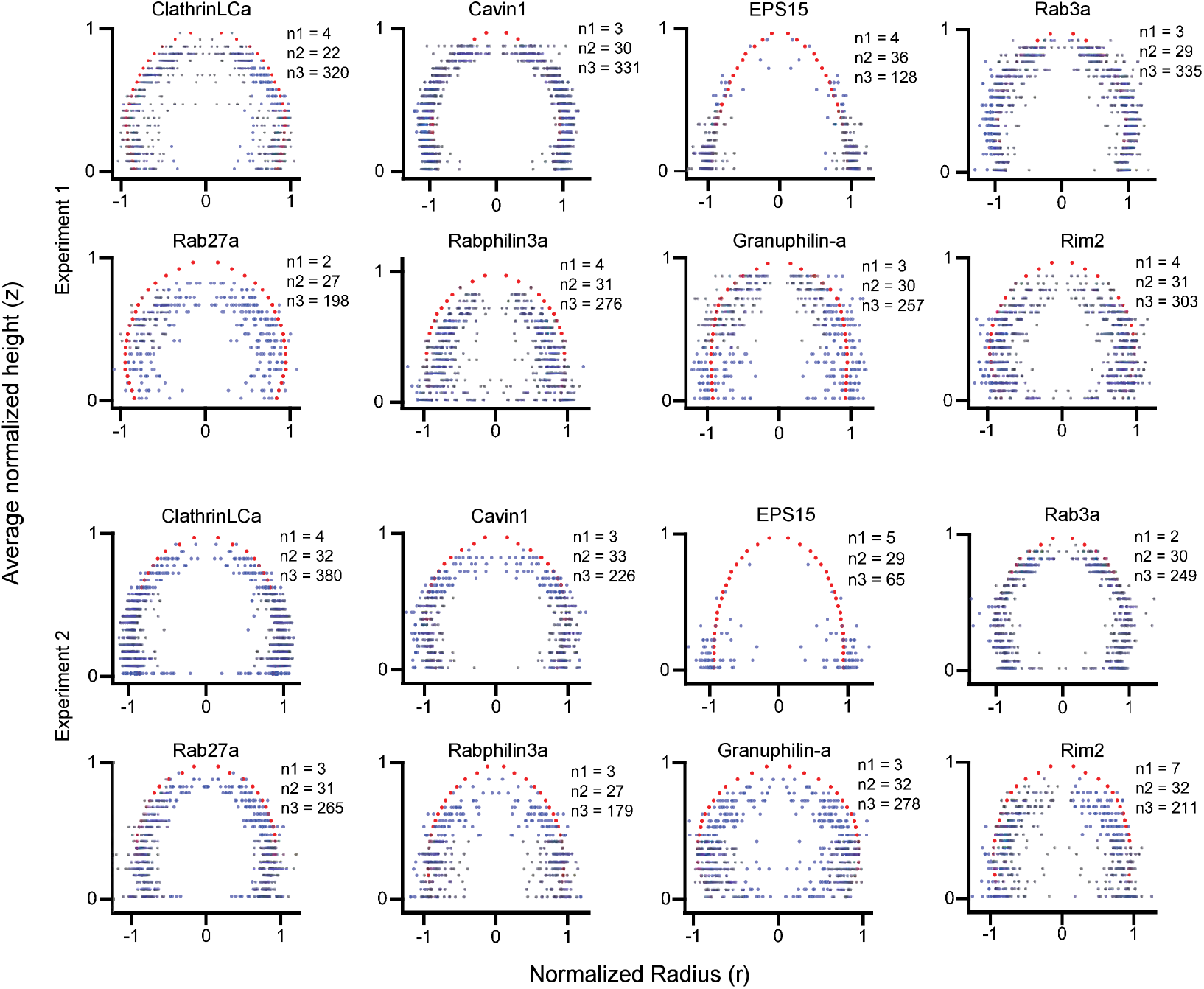
Vesicle profiles showing 3D distribution of gold on CCVs and DCVs in two independent experiments. Average vesicle membrane profile and gold nanoparticle distribution reproduced for the proteins Clathrin light chain A, Cavin1, EPS15, Rab27a, Rab3a, Rabphilin3a, Granuphilin-a, and Rim2 as seen in two independent experiments. The distribution is presented symmetrically. The notation nl, n2 and n3 refers to the number of tomograms, vesicles, and gold nanoparticles used in the data analysis, respectively.

**Supplementary Figure 5.**
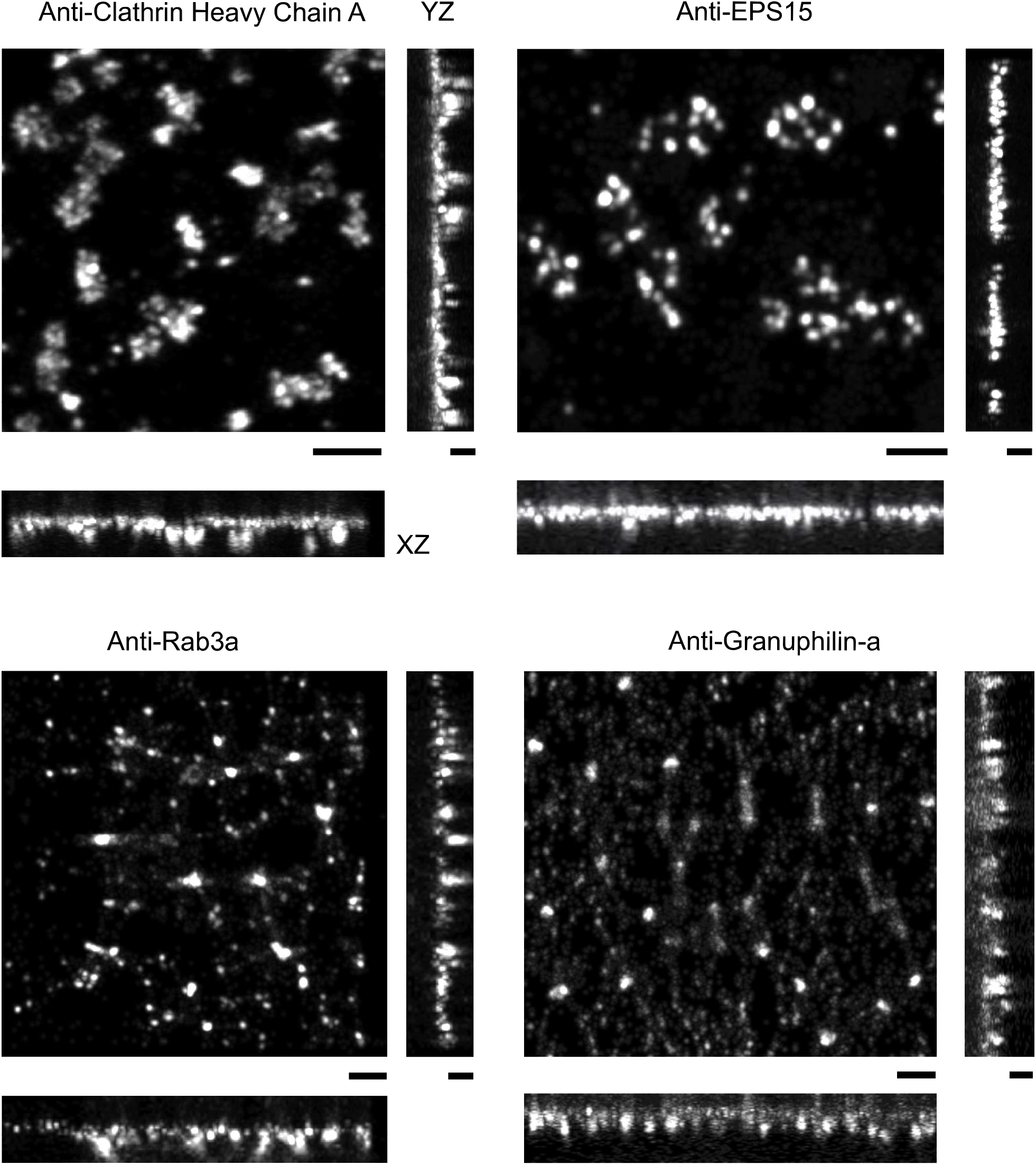
3D-STORM images showing distribution of endogenous proteins on CCVs and DCVs. Cells immunolabeled with anti-clathrin heavy chain (Hela), anti-EPS15 (Hela), anti-Rab3a (PC12), and anti-granuphilin-a (PC12) and fluorescently labeled with Alexa fluor 647 conjugated anti-rabbit or anti-mouse F(ab’)2 fragment. The figures show a maximum intensity projection of an XY (scale bar = 500 nm), XZ and YZ views (scale bar = 300 nm).

**Supplementary Figure 6.**
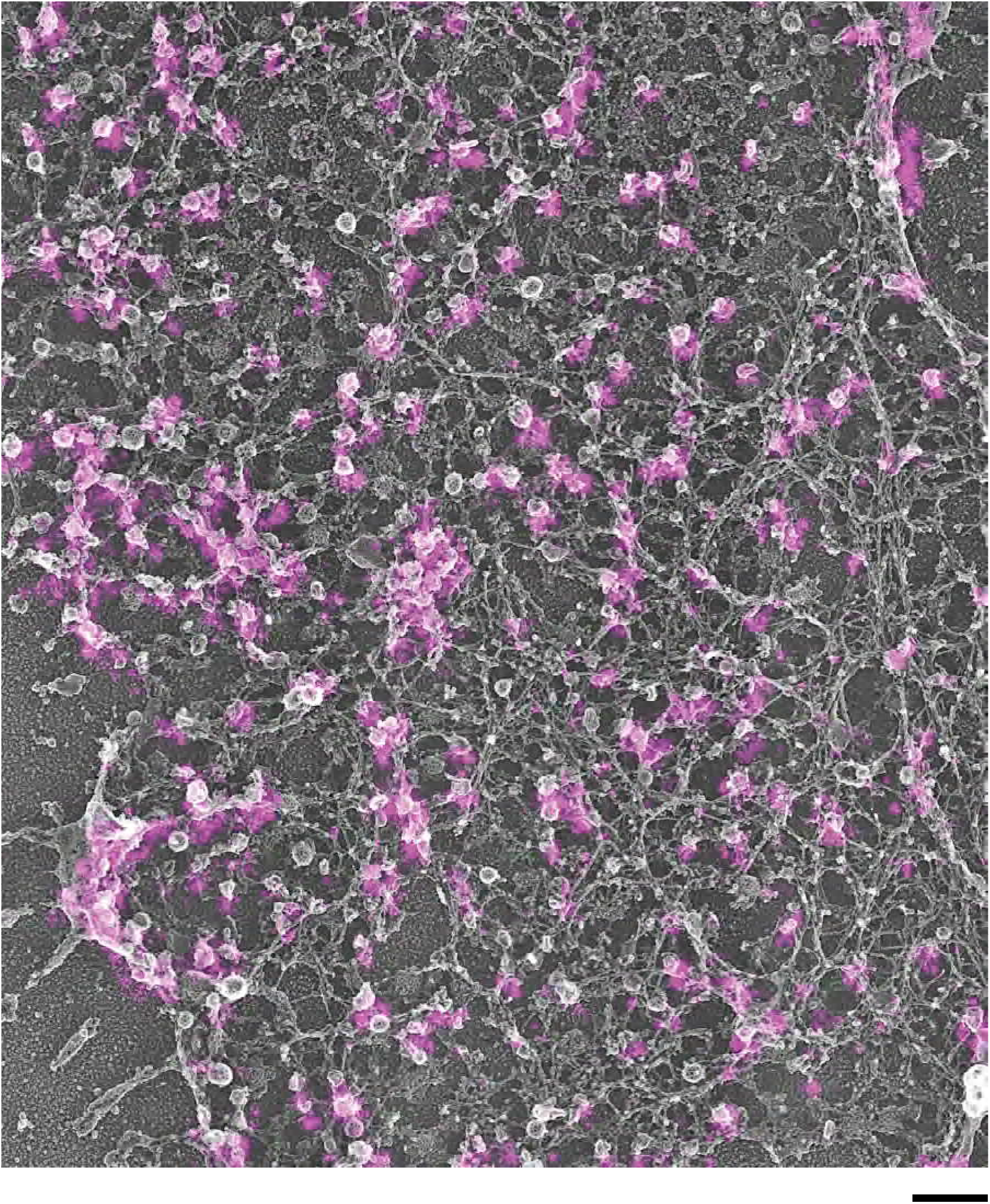
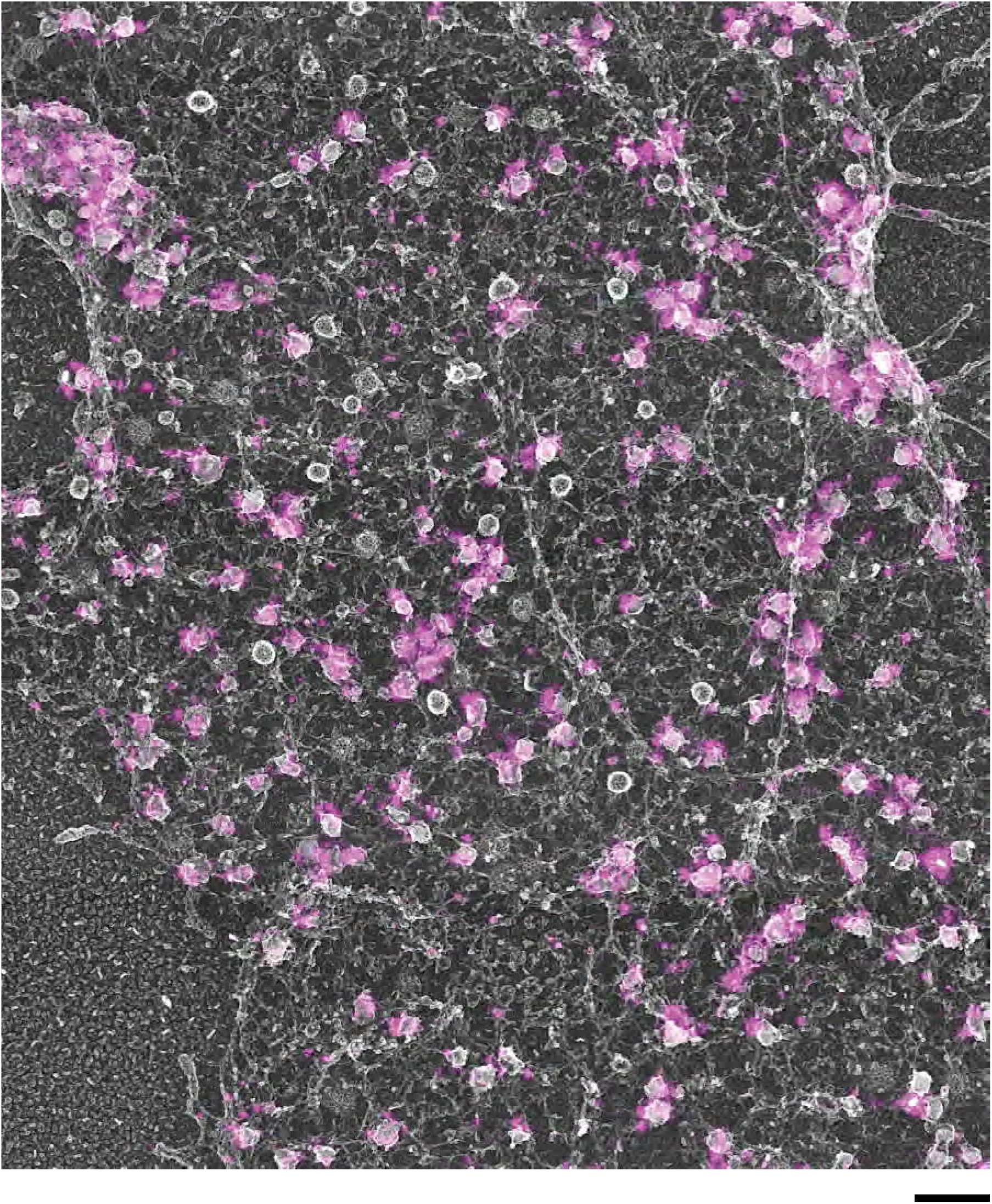
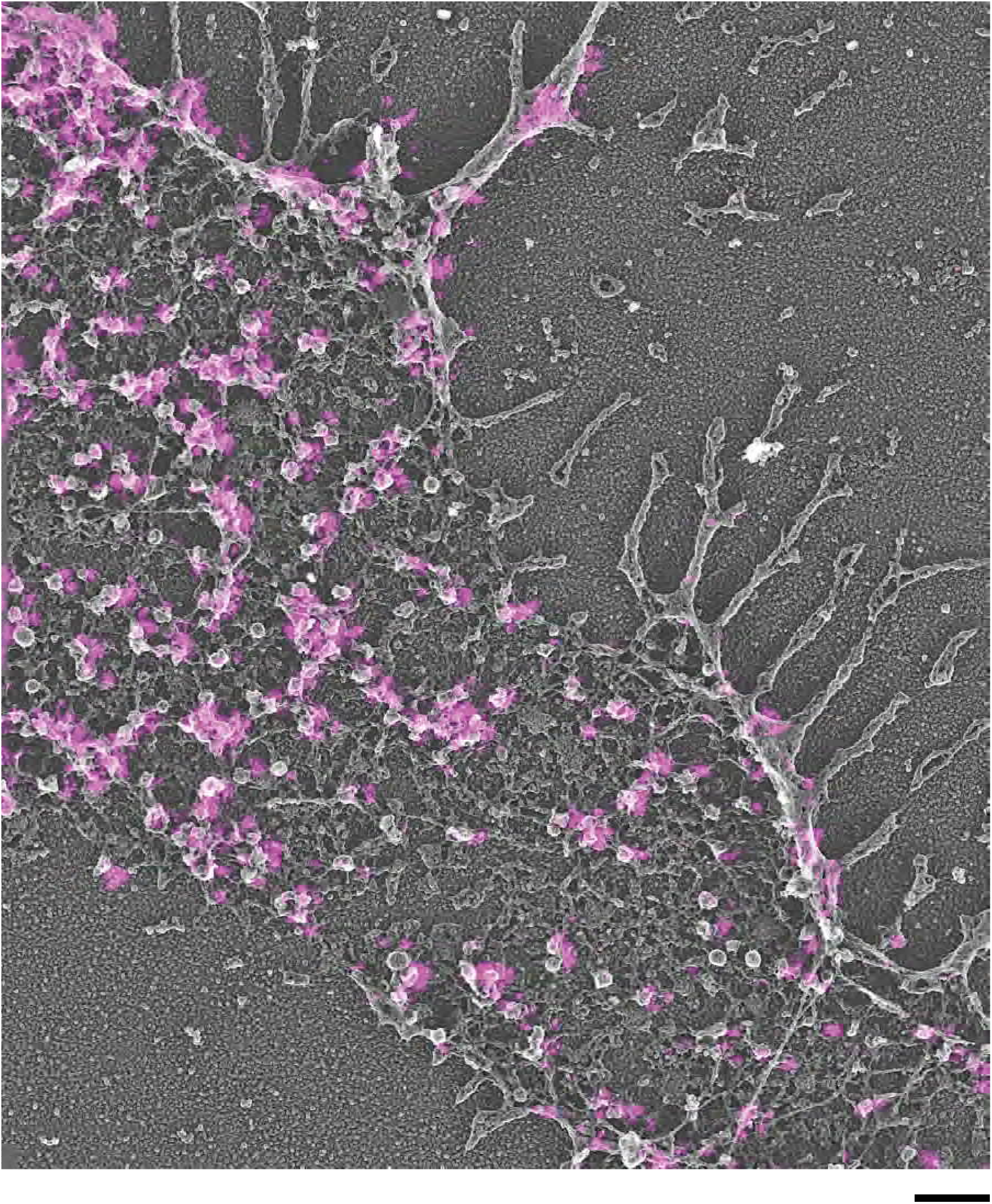
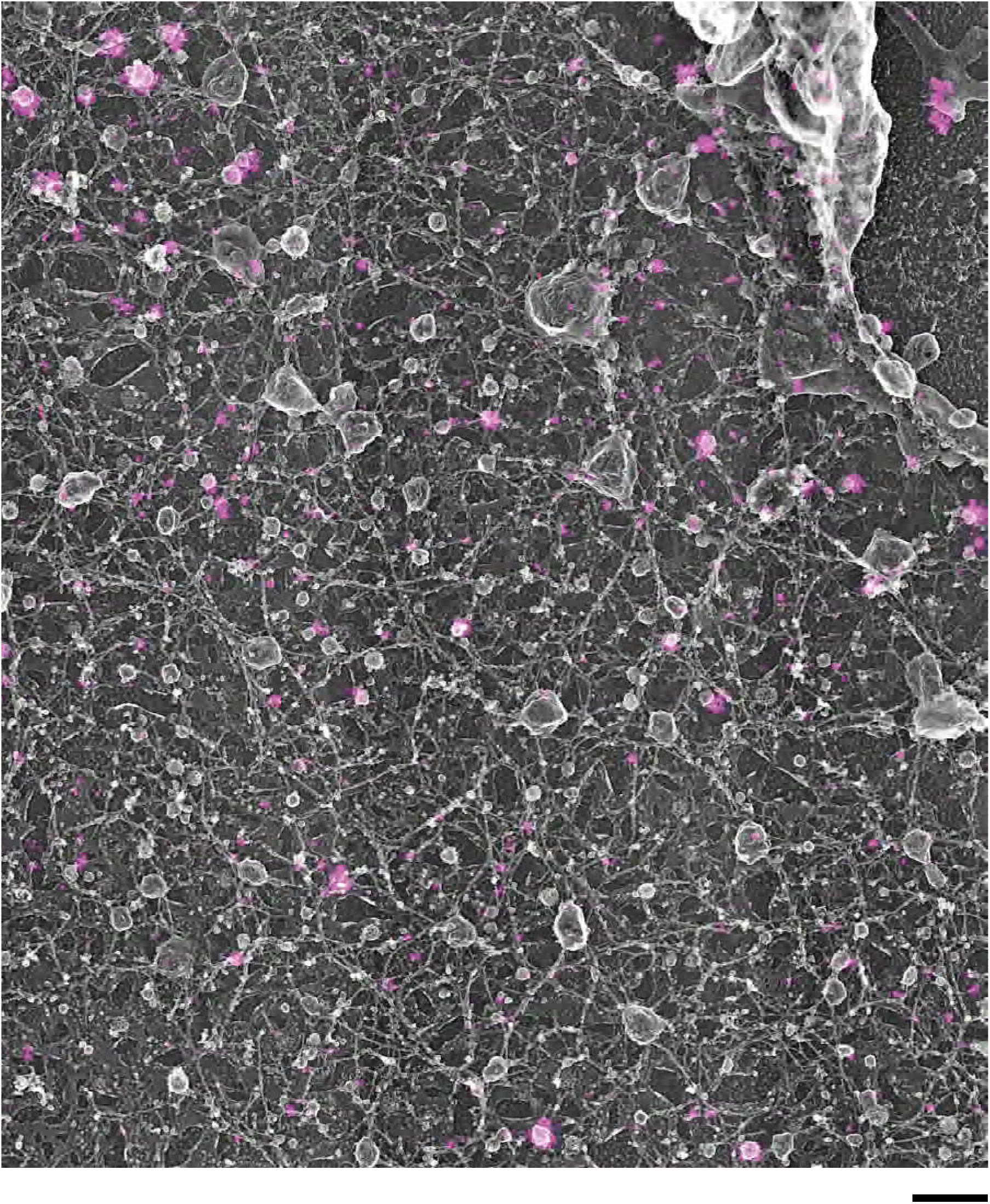
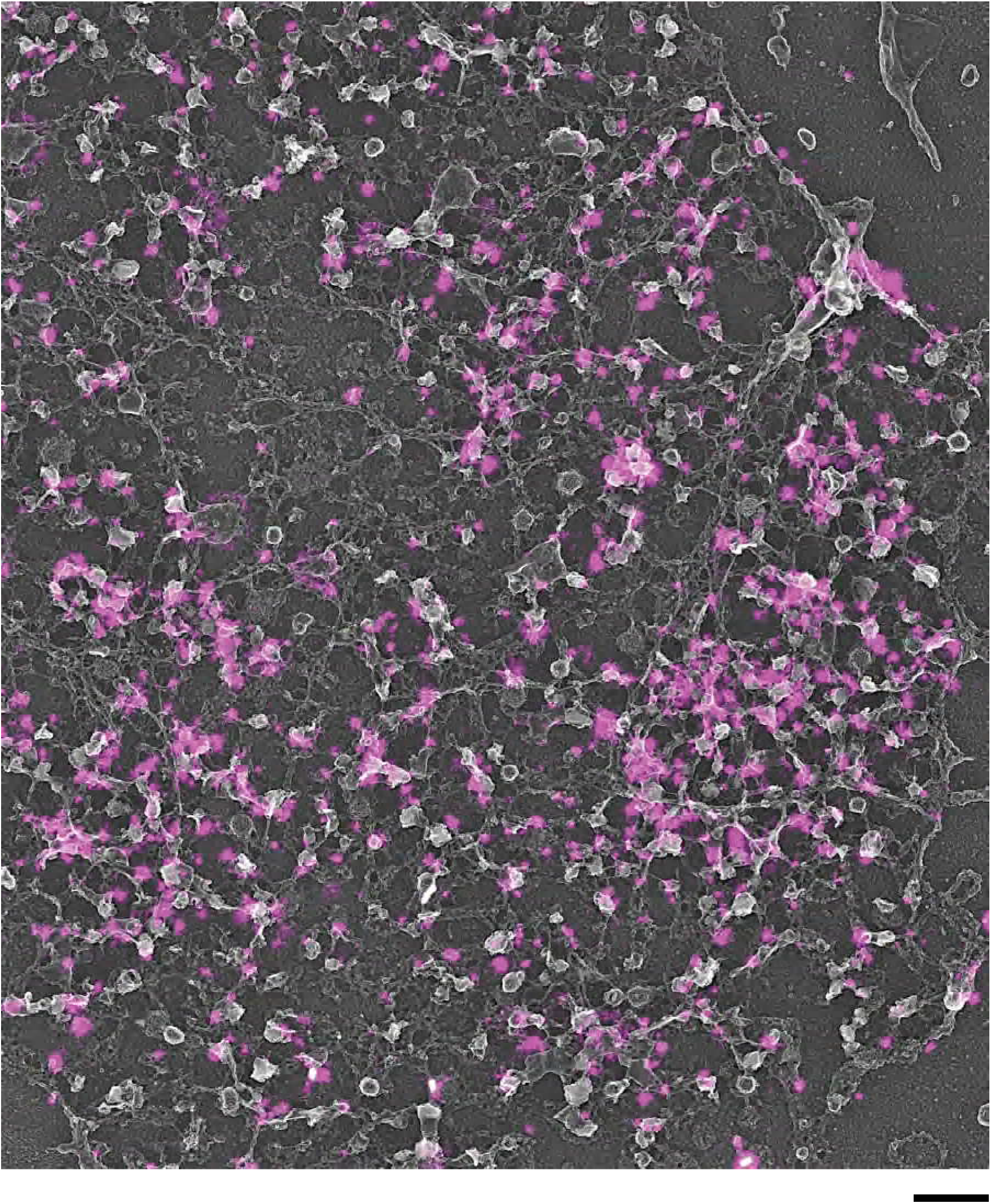
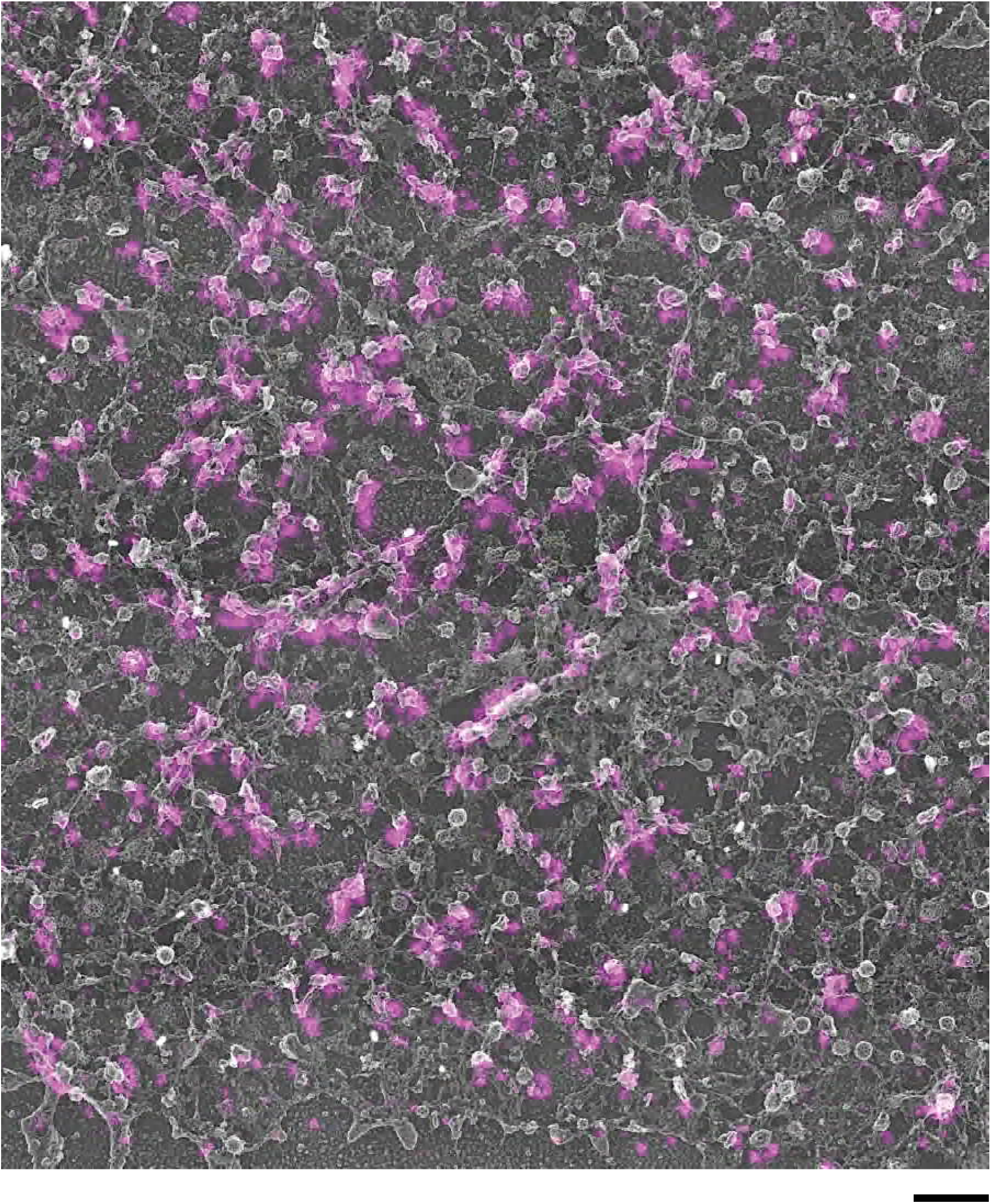
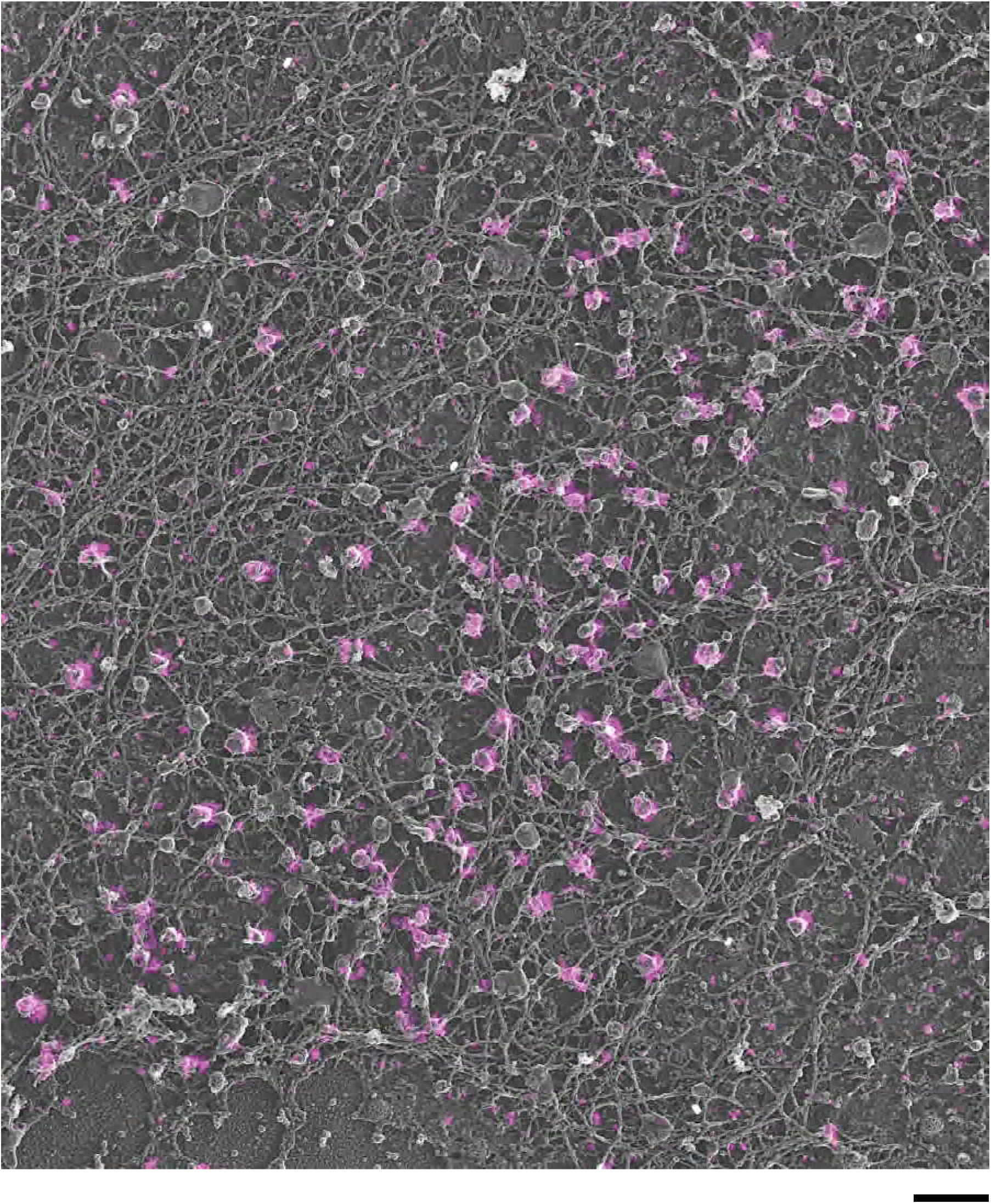
Original correlative STORM and TEM images of cells from which the cropped images in Figure 2 were derived. CLEM image for GFP-Rab3a, GFP-Rab27a, GFP-Rabphilin3a, GFP-Granuphilin-a, and immunolabeled Rim2, Rab3a, and Granuphilin-a in PC12 cells. Scale bars are 500 nm.

**Supplementary Figure 7.**
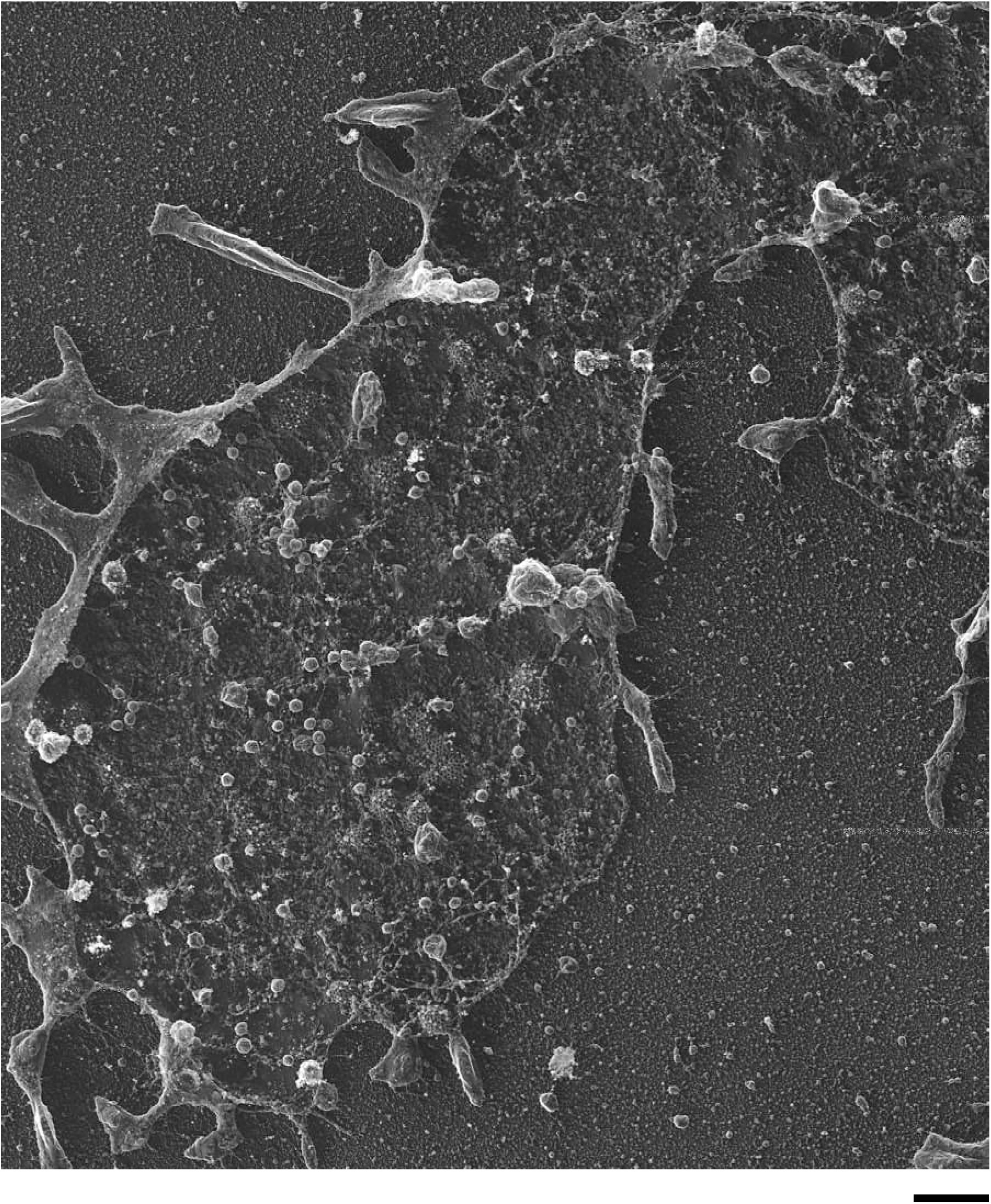
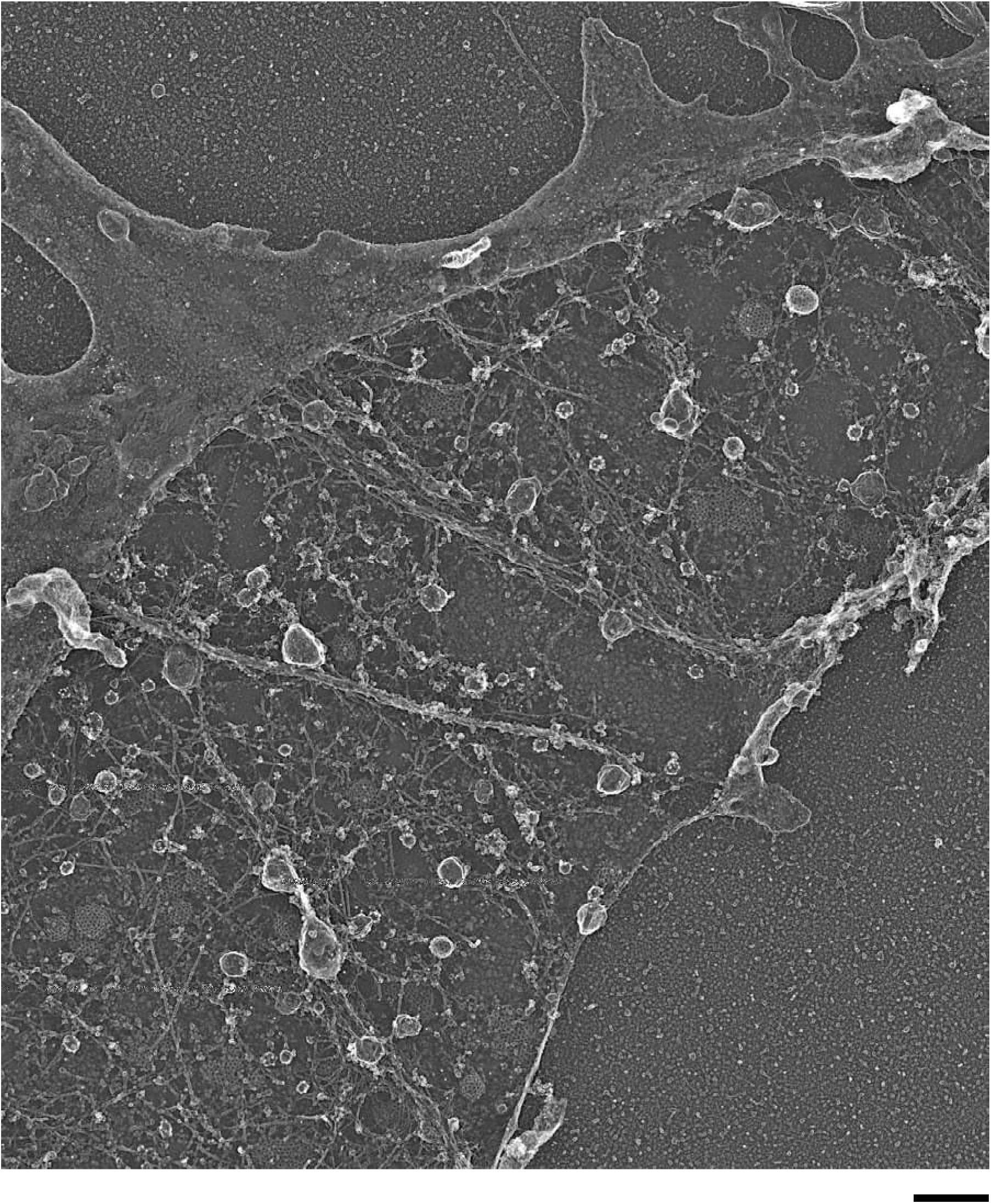
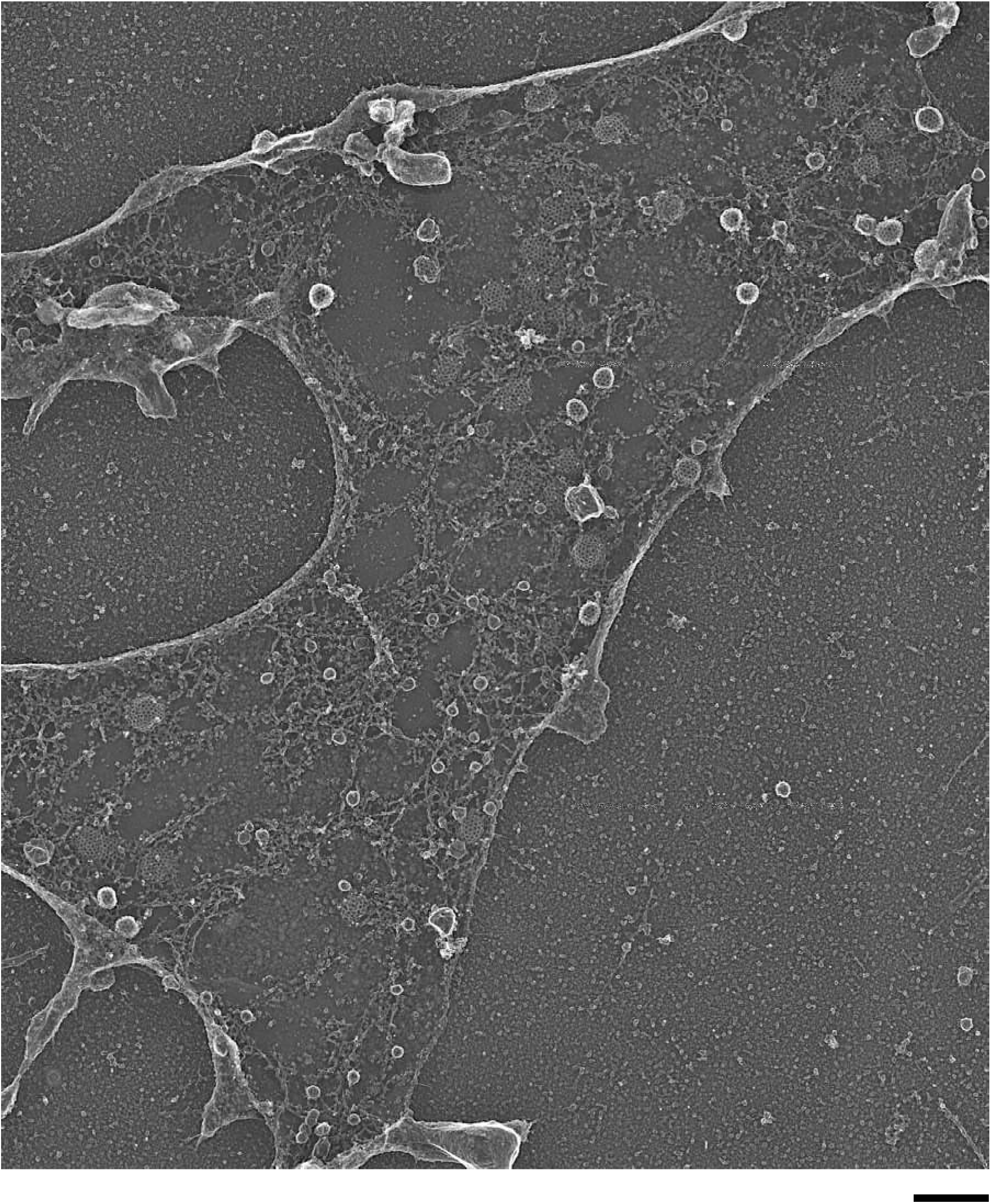
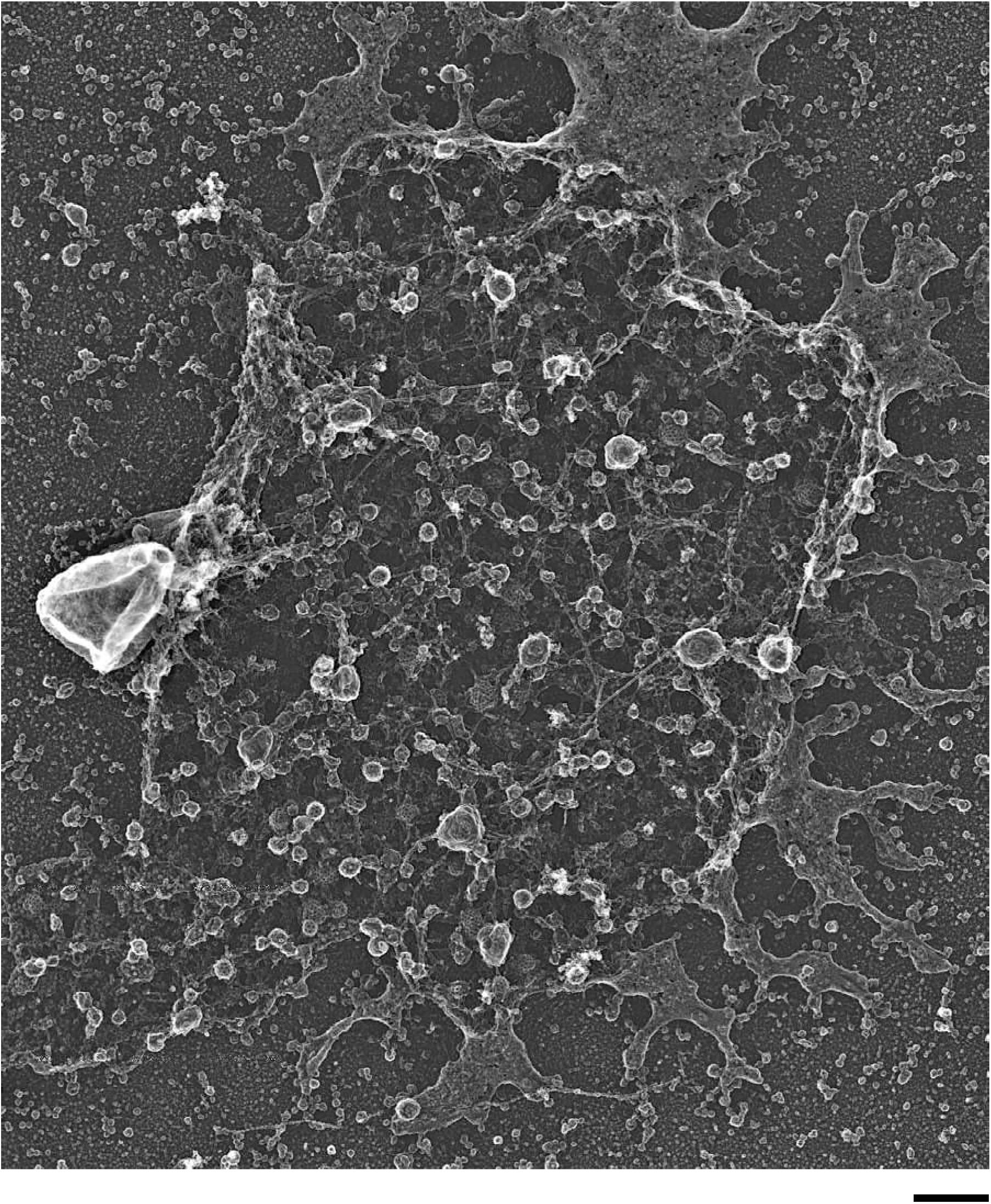
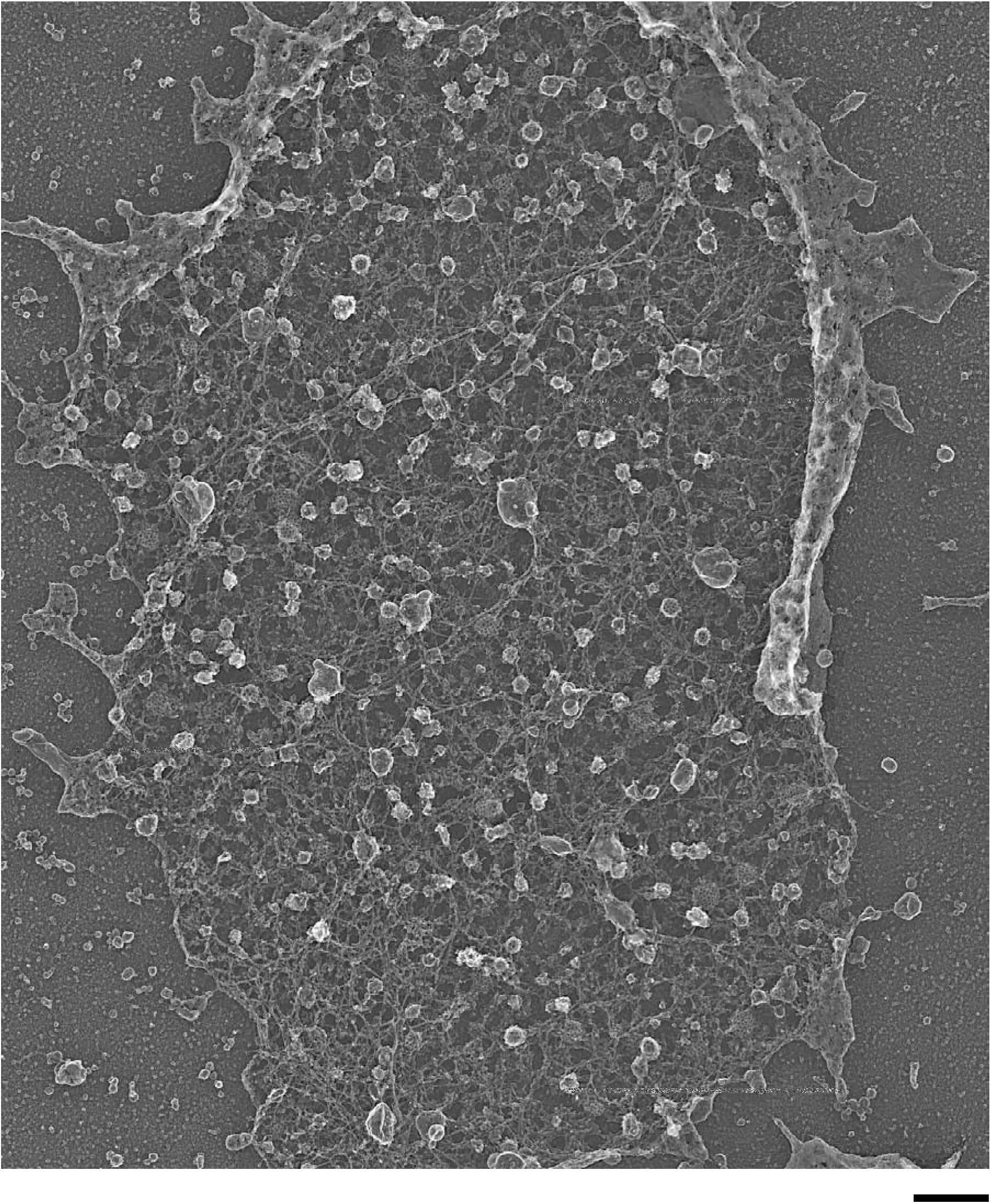
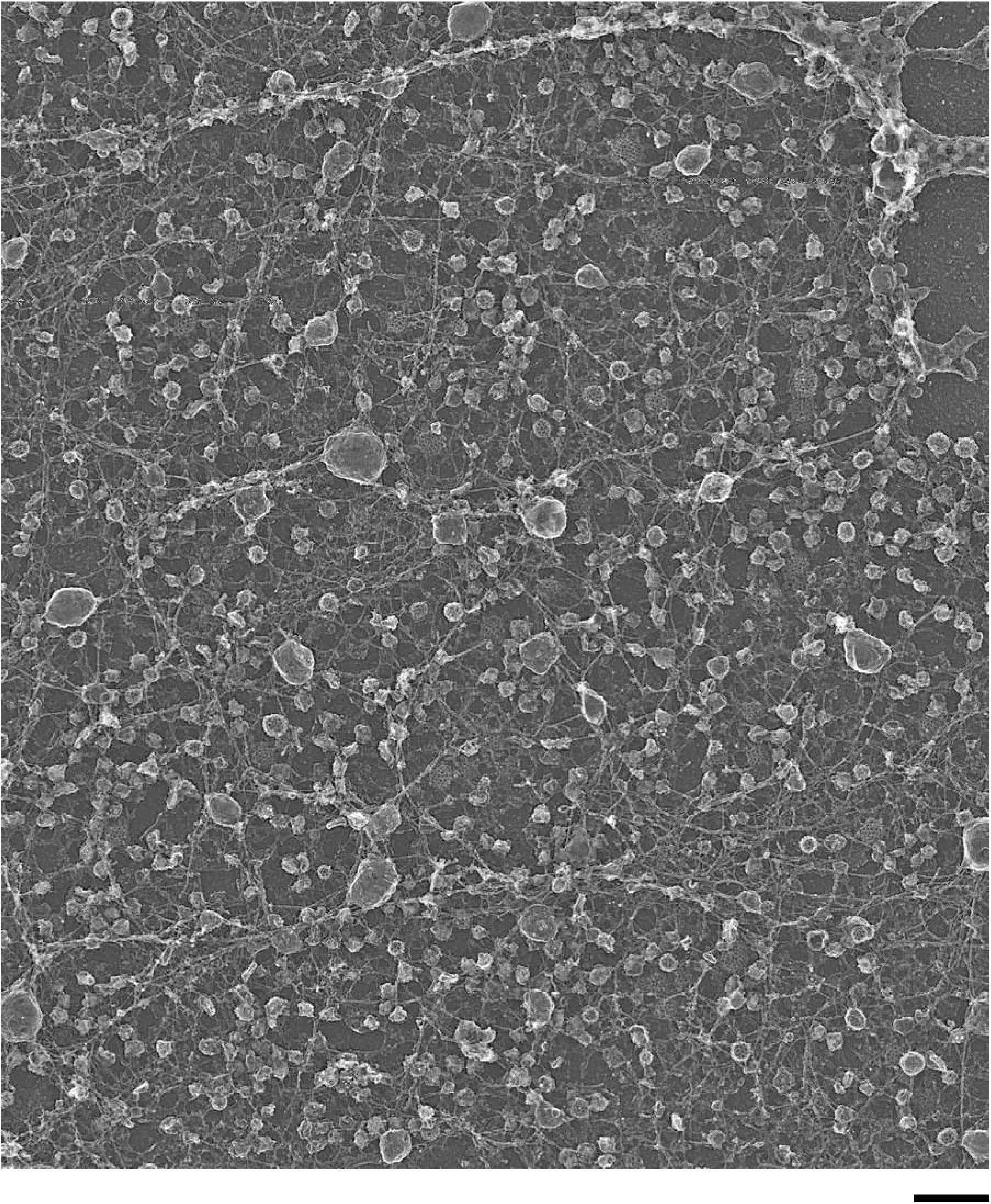
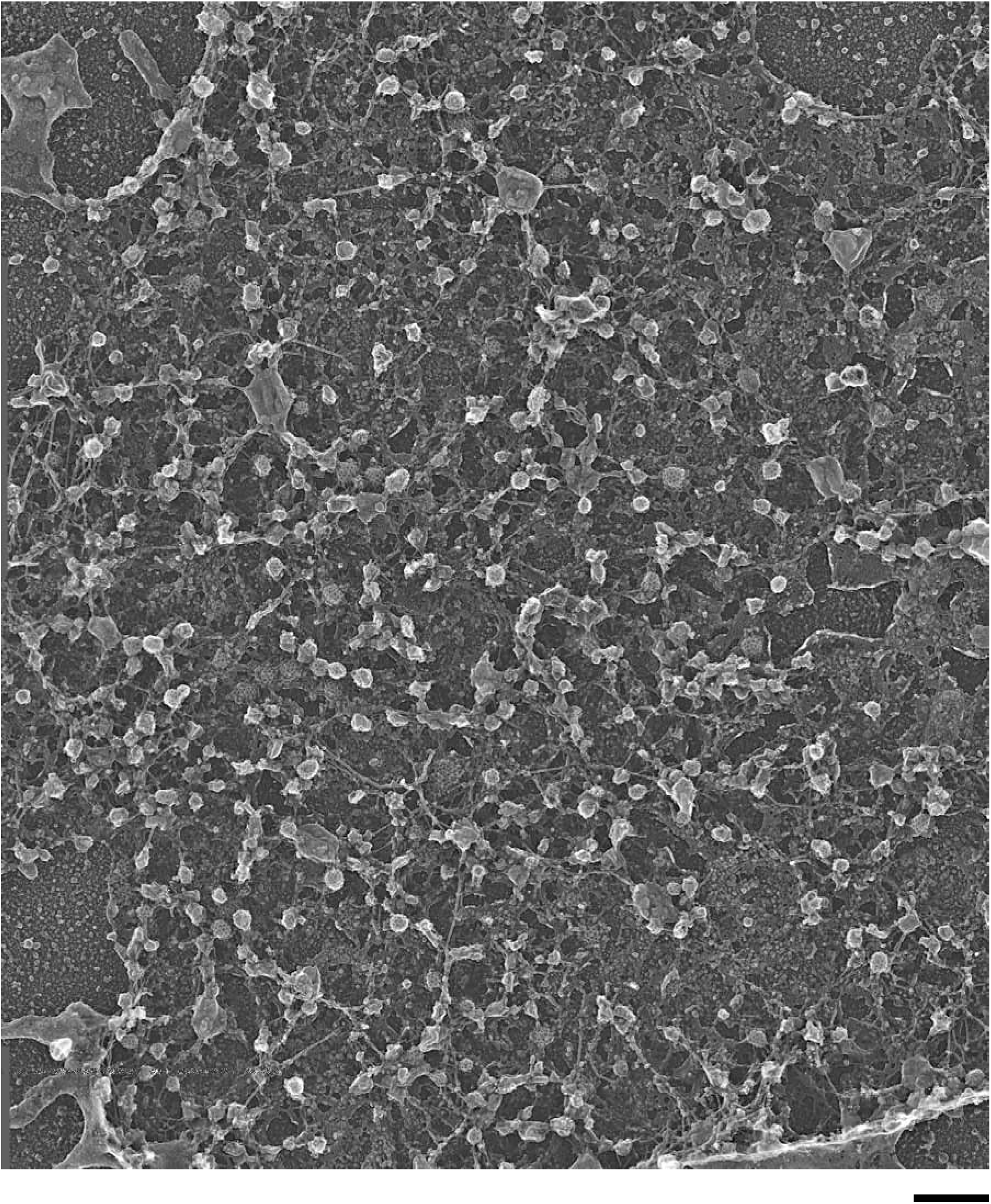
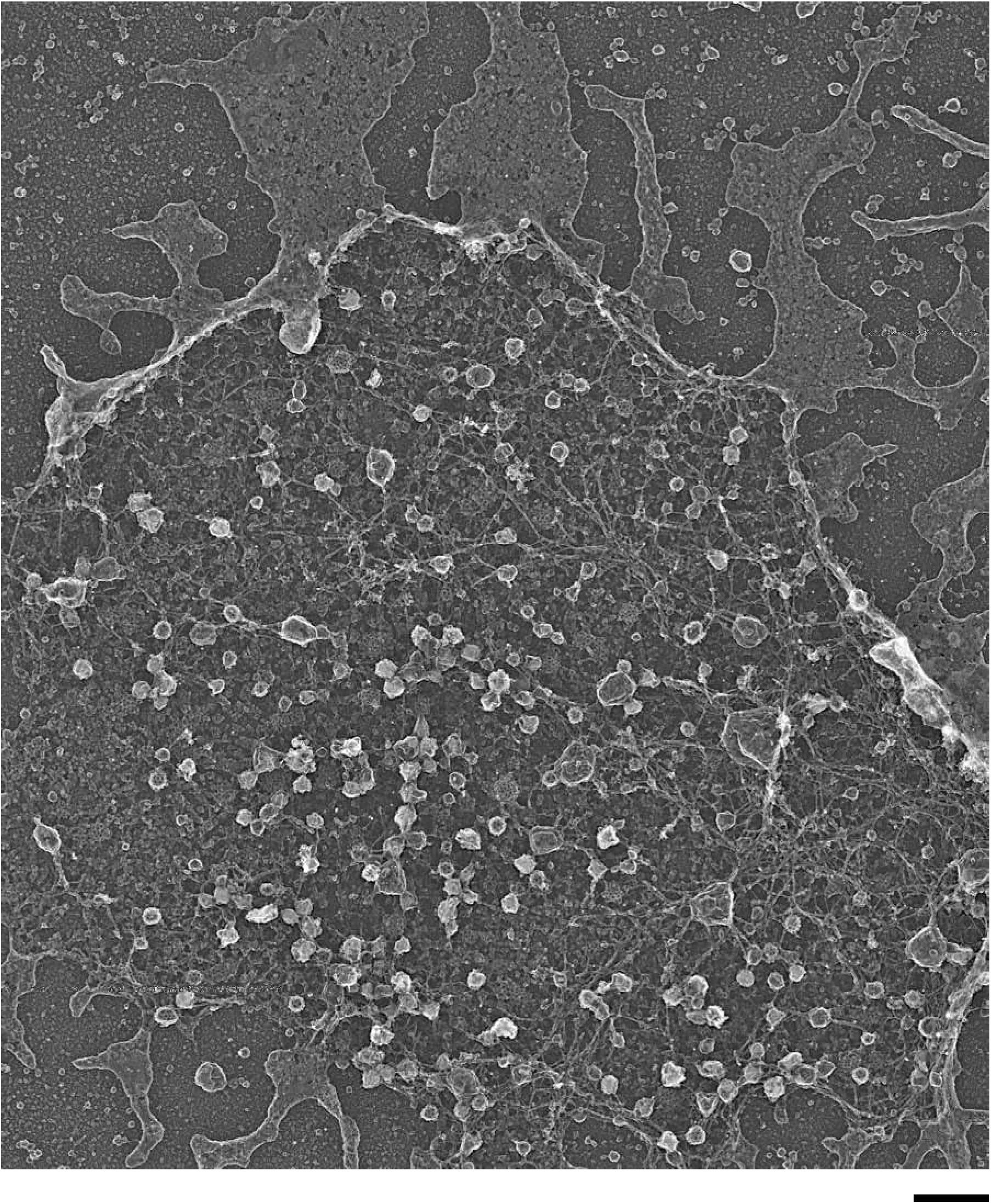
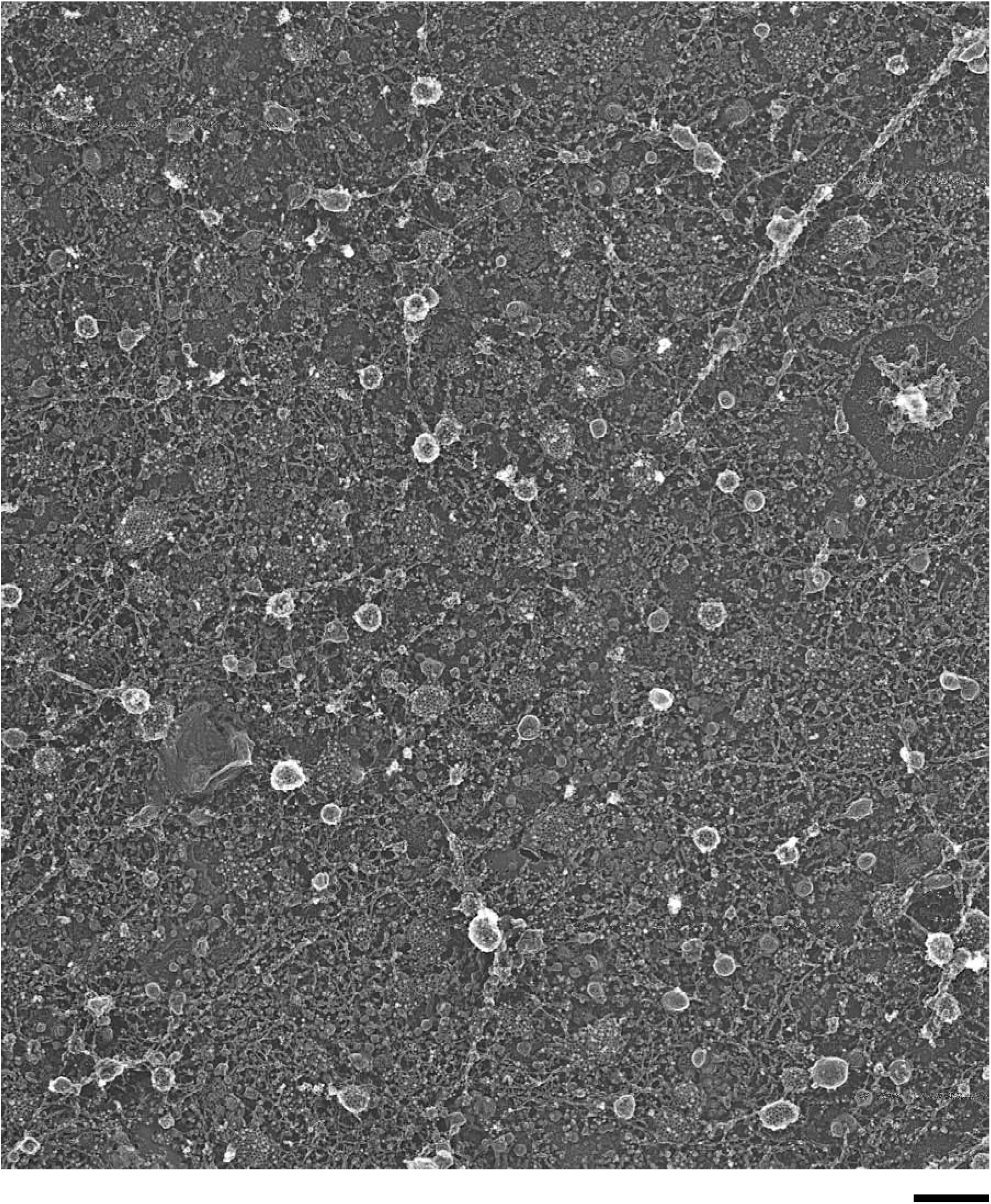
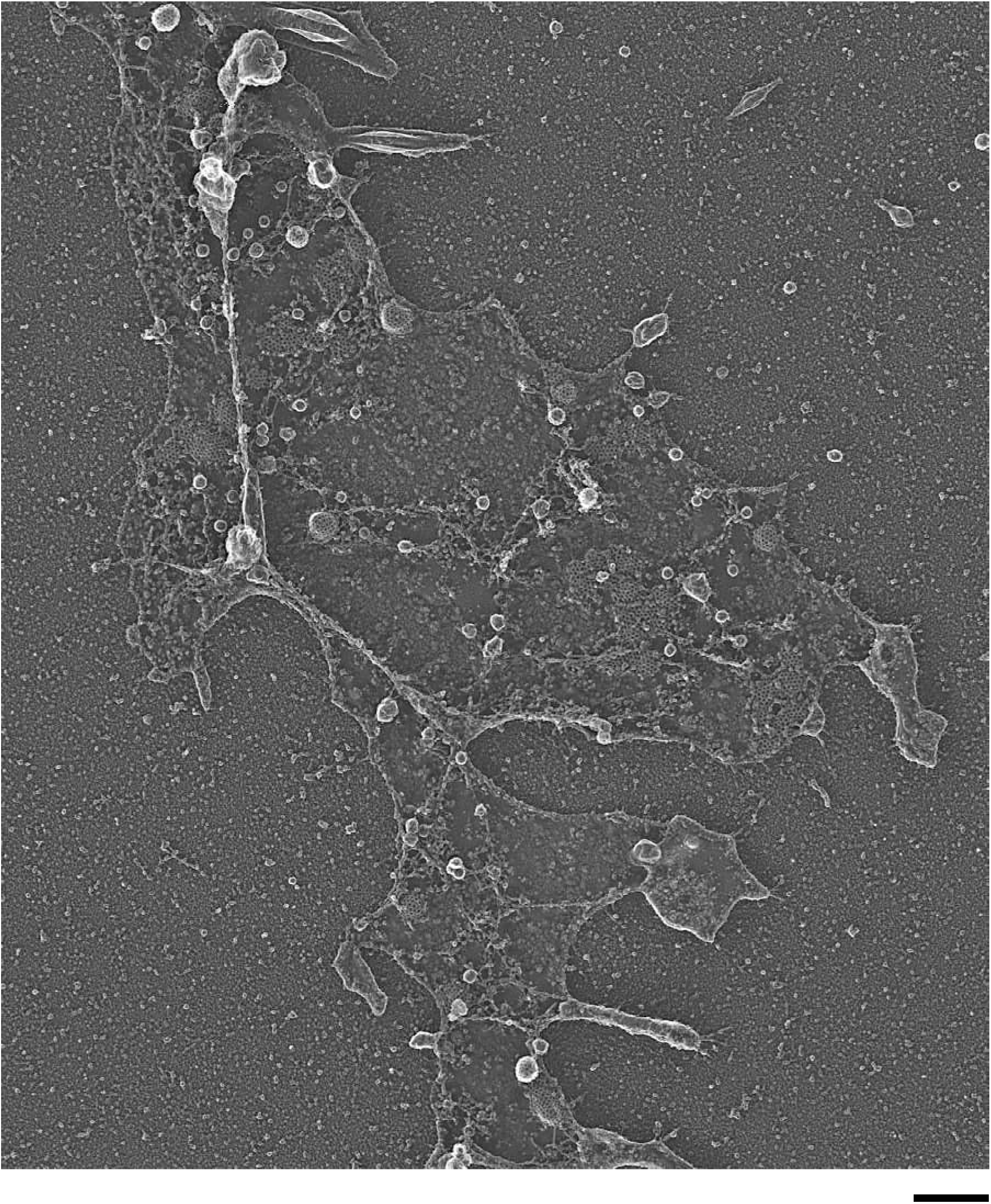
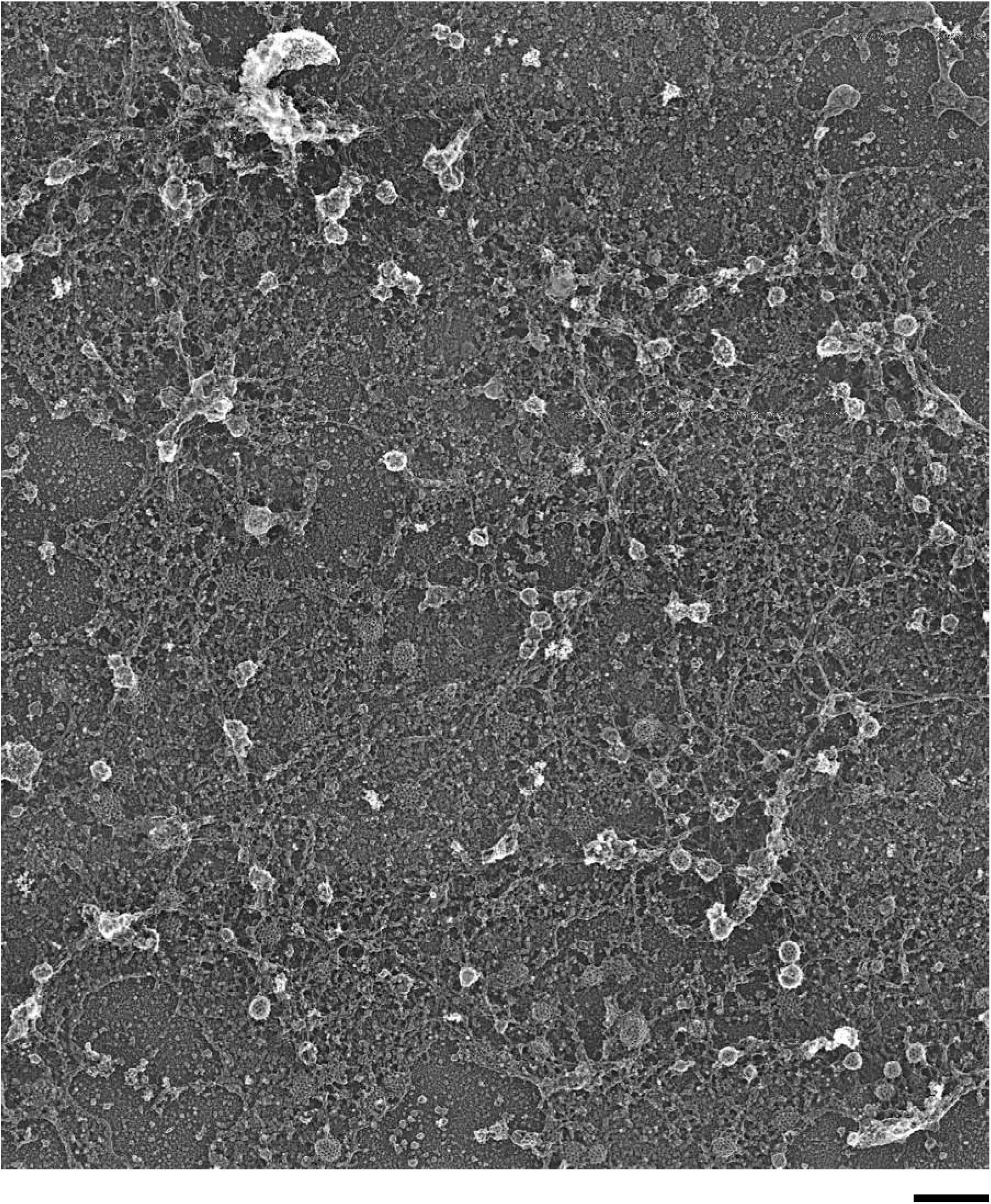
Original platinum replica TEM images of cells from which the cropped images in Figure 3 and 4 were derived. PREM images for cells transfected with His-GFP-clathrin light chain A, His-Cavin1-GFP, EPS15-GFP-His, His-GFP- Rab3a, His-GFP- Rab27a, His-GFP- Rabphilin3a, His-GFP- Granuphilin-a, and His-GFP-Rim2 and labeled with Ni-NTA-Au. PREM images for gold labeled U87-MG expressed with His-GFP-clathrin light chain A, Hela cells expressed with His-GFP-FCH02, and Insl cells expressed with His-GFP-Rab27a are also shown. Scale bars are 500 nm.

**Supplementary Table 1.** Table of samples used in the fluorescence profile analysis. The table shows the number of cells, coverslips, and dense core vesicles used in the analysis of dark GFP fusion proteins, Rab3a, Rab27a, Rabphilin3a, Granuphilin-a, and for immunolabeled Rab3a, Granuphilin-a, and Rim2. The number of vesicles analyzed for radial and edge profiles are listed separately.

|  | Cell # | Coverslip # | Vesicles # (radial) | Vesicles # (edge) |
| --- | --- | --- | --- | --- |
| GFP-Rab3a | 8 | 2 | 416 | 415 |
| GFP-Rab27a | 7 | 3 | 523 | 527 |
| GFP-Rabphilin3a | 10 | 3 | 635 | 624 |
| GFP-Granuphilina | 10 | 2 | 251 | 277 |
| Rab3a-Ig | 7 | 1 | 578 | 578 |
| Granuphilin-a-Ig | 2 | 1 | 216 | 208 |
| Rim2-Ig | 5 | 1 | 126 | 126 |
| Total | 49 | 13 | 2745 | 2755 |

**Supplementary Table 2.**
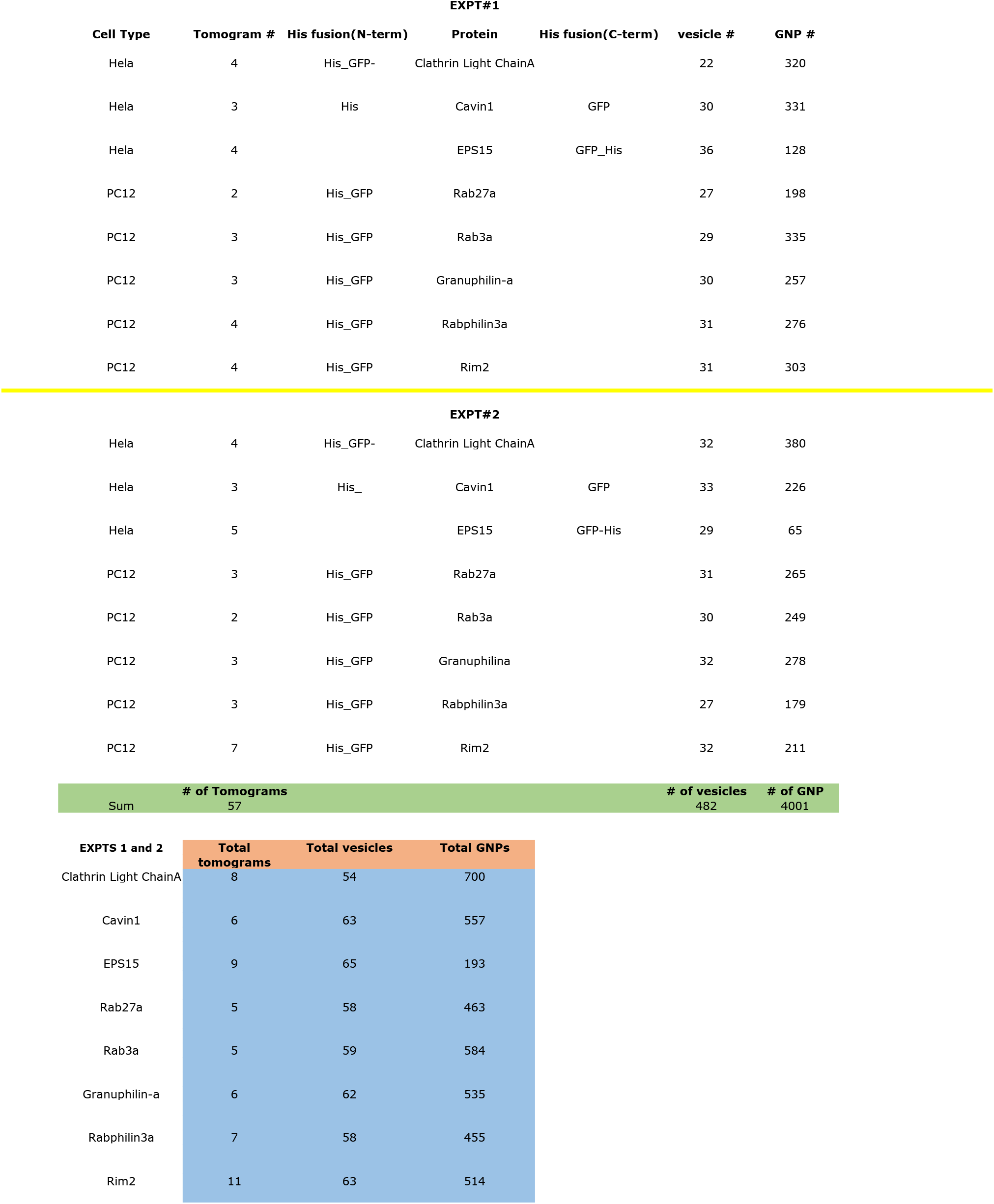
Table of number of cells, vesicles, and gold nanoparticles used in the tomogram analysis. The table shows types of cells used, number of tomograms, vesicles, and gold particles (GNPs) analyzed from two independent experiments and the combined info for His-GFP-CICa, Cavin-GFP-His, EPS15-GFP-His, His-GFP-Rab27a, His-GFP-Rab3a, His-GFP-Granuphilin-a, His-GFP-Rabphilin3a, His-GFP-Rim2 overexpressed cells.

**Supplementary Table 3.** Table of information about plasmids used in our study. The table includes the sequenced-confirmed plasmids used in CLEM experiments (red highlighted) and histidine based protein labeling with Ni-NTA-Au (all).

| Table of sequence confirmed plasmids |  |  |
| --- | --- | --- |
| Plasmid name | N-terminal protein | C-terminal protein |
| His-GFP-Clathrin | Histidine & EGFP | Mus musculus clathrin, light polypeptide (Lca) |
| His-Cavin-1-GFP | Histidine & Mus musculus caveolae associated 1 (Cavin1) | EGFP Cloning vector ENCODE_CRISPR_GFP<br>GFP tag fusion protein gene |
| EPS15-GFP-His | Homo sapiens epidermal growth factor receptor pathway substrate 15<br>(EPS15) | monomeric EGFP (A206K) |
| His-GFP-Rab27a | Histidine & EGFP | Homo sapiens RAB27A, member RAS<br>oncogene family (Rab27A) |
| His-GFP-Rab3a | Histidine & EGFP | Mus musculus RAB3A, member RAS<br>oncogene family (Rab3a) |
| His-GFP-Rabphilin | Histidine & EGFP | Bos taurus rabphilin 3A homolog |
| His-GFP-Granuphilin | Histidine & EGFP | Homo sapiens synaptotagmin-like 4 |
| His-GFP-Rim2 | Histidine & EGFP | Rattus norvegicus RIM2 |

**Supplementary Video 1.** Tomogram of a Hela cell labeled with Ni-NTA-Au against His-GFP-clathrin light chain A. The tomogram is viewed in the XY plane through Z slices. Each voxel is 2.3 nm in all dimensions. The tomogram is 2.4 mm in width and 1.9 mm in height.

**Supplementary Video 2.** Tomogram of a Hela cell labeled with Ni-NTA-Au against His-cavin1-GFP. The tomogram is viewed in the XY plane through Z slices. Each voxel is 2.3 nm in all dimensions. The tomogram is 2.4 mm in width and 1.9 mm in height.

**Supplementary Video 3.** Tomogram of a Hela cell labeled with Ni-NTA-Au against EPS15-GFP-His. The tomogram is viewed in the XY plane through Z slices. Each voxel is 2.3 nm in all dimensions. The tomogram is 2.4 mm in width and 1.9 mm in height.

**Supplementary Video 4.** Tomogram of a PC12 cell labeled with Ni-NTA-Au against His-GFP-Rab27a. The tomogram is viewed in the XY plane through Z slices. Each voxel is 2.3 nm in all dimensions. The tomogram is 2.3 mm in width and 1.9 mm in height.

**Supplementary Video 5.** Tomogram of a PC12 cell labeled with Ni-NTA-Au against His-GFP-Rab3a. The tomogram is viewed in the XY plane through Z slices. Each voxel is 2.3 nm in all dimensions. The tomogram is 1.6 mm in width and 1.8 mm in height.

**Supplementary Video 6.** Tomogram of a PC12 cell labeled with Ni-NTA-Au against His-GFP-Granuphilin-a. The tomogram is viewed in the XY plane through Z slices. Each voxel is 2.3 nm in all dimensions. The tomogram is 2.3 mm in width and 1.9 mm in height.

**Supplementary Video 7.** Tomogram of a PC12 cell labeled with Ni-NTA-Au against His-GFP-Rabphilin3a. The tomogram is viewed in the XY plane through Z slices. Each voxel is 2.3 nm in all dimensions. The tomogram is 2.3 mm in width and 1.9 mm in height.

**Supplementary Video 8.** Tomogram of a PC12 cell labeled with Ni-NTA-Au against His-GFP-Rim2. The tomogram is viewed in the XY plane through Z slices. Each voxel is 2.3 nm in all dimensions. The tomogram is 2.9 mm in width and 1.9 mm in height.

